# Robust Classification of Protein Variation Using Structural Modeling and Large-Scale Data Integration

**DOI:** 10.1101/029041

**Authors:** Evan H. Baugh, Riley Simmons-Edler, Christian L. Müller, Rebecca F. Alford, Natalia Volfovsky, Alex E. Lash, Richard Bonneau

**Affiliations:** Department of Biology, New York University, NY, NY 10003; Computer Science Department, New York University, NY, NY 10003; New York University Center for Genomics and Systems Biology, NY, NY 10003; Simons Foundation, NY, NY 10010; Simons Center for Data Analysis, Simons Foundation, NY, NY 10010; Carnegie Mellon University Department of Chemistry, 5000 Forbes Ave, Pittsburgh, PA, 15289; Commack High School, Commack NY, 11725

## Abstract

Existing methods for interpreting protein variation focus on annotating mutation pathogenicity rather than detailed interpretation of variant deleteriousness and frequently use only sequence-based or structure-based information. We present VIPUR, a computational framework that seamlessly integrates sequence analysis and structural modeling (using the Rosetta protein modeling suite) to identify and interpret deleterious protein variants. To train VIPUR, we collected 9,477 protein variants with known effects on protein function from multiple organisms and curated structural models for each variant from crystal structures and homology models. VIPUR can be applied to mutations in any organism’s proteome with improved generalized accuracy (AUROC .83) and interpretability (AUPR .87) compared to other methods. We demonstrate that VIPUR‘s predictions of deleteriousness match the biological phenotypes in ClinVar and provide a clear ranking of prediction confidence. We use VIPUR to interpret known mutations associated with inflammation and diabetes, demonstrating the structural diversity of disrupted functional sites and improved interpretation of mutations associated with human diseases. Lastly we demonstrate VIPUR‘s ability to highlight candidate genes associated with human diseases by applying VIPUR to de novo variants associated with autism spectrum disorders.

## INTRODUCTION

High-throughput sequencing technologies and new computational techniques for analyzing population genetics data are rapidly improving our understanding of disease susceptibility in humans(29, 32, 51) and adaptation in a wide variety of organisms, including crop species and pathogens(10, 41, 50). These studies often discover nonsynonymous variation with large effects as even a single amino acid change can disrupt the folding, catalytic activity, and physical interactions of proteins(11, 27). Current estimates predict that every human genome contains 10,000-11,000 nonsynonymous variations(46, 48) and, while we cannot currently characterize all this diversity experimentally, many variants that alter protein function can be identified computationally from destabilization of structural models or amino acid conservation(8, 10, 47). Methods for annotating variant effects in genome-wide association studies and exome sequencing studies, such as PolyPhen2(1), CADD(22), PROVEAN(7), and SIFT(35), use conservation and other sequence-based features to identify damaging variants but cannot predict the effect these variants have on protein function. Recent studies of *de novo* variants(9, 12, 36, 38) have demonstrated the power of these methods but also the need for additional information(10), such as physical models from the Protein Data Bank(4), to identify causal variants in disease association studies.

Most methods for annotating coding variants attempt to predict variant deleteriousness in the context of the whole orangism (where deleteriousness is defined as the tendency for a variant to reduce organismal fitness, to express an altered phenotype, or to exhibit an association with a disease condition)(22). Deleteriousness, when defined in terms of fitness or phenotypic effects, is difficult to measure directly but underlies patterns of conservation, molecular functionality, and disease pathogenicity. Variant annotations in several databases are often limited to discrete labels such as deleterious or neutral. Definitions based on deleteriousness are often confused with definitions of pathogenicity used to curate training and benchmarking on datasets. The annotations predicted by current coding variant annotation methods for these reasons have diverse implications. For example, SIFT segregates “tolerant” from “intolerant” variants(35) while PolyPhen2 identifies “possibly damaging” and “probably damaging” effects(1). CADD predicts deleteriousness by distinguishing fixed from simulated variation and relies on the predictions of other methods including both SIFT and PolyPhen2(22). Each of these methods predicts a label that is designed to correlate with variant deleteriousness and is used to prioritize causal pathogenic variants from large genomic datasets(10). Deleteriousness can be approximated with measures of conservation and molecular functionality but available data on both protein sequence variation and structural energetics are rarely combined(6, 40, 43). Selection against deleterious variants can be detected by analysis of conservation and other alignment-based methods, although these metrics may not apply to *de novo* mutations. Alternatively, several studies have aimed to model the biophysical characteristics of mutations, such as stability, enzymatic function, and the *pK_a_* of key residues. Protein structure models of mutations can be used to indicate disruption of active sites and destabilization of the folded protein (6, 11, 20, 45) using tools like Rosetta (20, 25) and FoldX (45). Here we aim to provide a measure of deleteriousness centered on individual proteins with deleterious defined as disrupted protein function (disrupted stability, active site, interface, or folding). Our method aims to use conservation and structural analyses to better predict protein-centered deleteriousness.

We present **VIPUR** (*vIpә(r)*, **V**ariant **I**nterpretation and **P**rediction **U**sing **R**osetta), a computational framework capable of identifying, ranking, and interpreting deleterious protein variants in different species. To make VIPUR applicable across multiple species, we curated **VTS** (the VIPUR **T**raining **S**et), a novel collection of 9,477 annotated variants from >360 species containing both natural variations and experimental mutations. Variant annotations were carefully curated, restricting **VTS** to **<u>deleterious</u>** variants which directly disrupt protein molecular function or are functionally **<u>neutral</u>**, rather than “pathogenicity” or “intolerance”. We obtained structural models for these proteins from solved crystal structures and comparative modeling initiatives, such as ModBase(39), taking advantage of reliable homology models freely available for most human proteins. Structural analysis is performed using Rosetta to rigorously sample variant protein conformations, properly accommodating the variant amino acid by moving the protein backbone(20, 40, 49). We combine sequence-based and structure-based features in a sparse logistic regression framework, leading to a classifier that accurately ranks deleterious variants, with ≥90% precision on the highest scoring 3,800 variants (40% of variants classified) and 0.872 Area Under the Precision-Recall curve (AUPR). In addition to classification and ranking, VIPUR uses structural analysis to provide a detailed prediction of each variant’s physical effect, automatically reporting disruption of hydrogen bonding, side-chain packing, and backbone stability.

VIPUR deleterious predictions do not guarantee the presence of a disease phenotype. Nonetheless, distributions of VIPUR **<u>deleterious</u>** scores match the expectation for known **pathogenic** and **benign** variant phenotypes in ClinVar(24) while PolyPhen2 produces many false positives that overshadow true positives when applied to variants with uncertain effects. We apply VIPUR to a small set of variants (388) in proteins associated with inflammation and diabetes mellitus to identify deleterious variants improperly annotated by sequence-based methods and demonstrate the clarity of VIPUR predictions. We demonstrate the ability of VIPUR to identify and rank potentially causal variants in the *de novo* missense mutations of the Simons Simplex Collection(19, 37, 42) and compare to other variant annotation methods (2,226 missense variants).VIPUR **<u>deleterious</u>** predictions demonstrate a clear enrichment for mutations found in children with autism that is unmatched by current variant annotation methods and highlights a small set of extremely confident candidate genes for future investigation.

## MATERIALS AND METHODS

### Generating a Deleterious Protein Variant Benchmark

Existing datasets for the training and benchmarking of protein variant annotation methods are frequently restricted in scope, focusing on disease-associated variants(1, 7, 15, 34). Methods that model protein structures are similarly restricted, validating on *in vitro* experimental characterization of variants produced by mutagenesis(6, 45). We want VIPUR to predict variant deleteriousness and generalize to both natural variants *and* mutagenesis variants. We collected and curated missense variants from multiple experimental sources and prepared structural models from different databases to ensure VIPUR is benchmarked on diverse protein structures (see Supplementary Figure S1). Protein variants from HumDiv(1) and UniProt(2) with clear ‘**<u>deleterious</u>**’ or ‘**<u>neutral</u>**’ effects were mapped onto crystallographic and comparative models of the protein macromolecules from the Protein Data Bank(4), ModBase(39), and SwissModel(44). Our **<u>deleterious</u>** and **<u>neutral</u>** labels are restricted to variants with direct evidence of protein disruption, avoiding the assumptions that all disease-associated variants are necessarily deleterious(1) or that all unannotated variants are necessarily neutral(7). This training set, **VTS**, includes 9,477 variants (5,740 deleterious, 3737 neutral, 1.54 label ratio) and curated structural models (2,637 models in 2,444 proteins), available at <u>https://osf.io/bd2h4</u>. Each variant is characterized by 106 sequence and structure features (see below). **VTS** comprises 5,901 human variants, 1,635 variants in other Eukaryotic proteins, 1,725 in Prokaryotic proteins, 122 Archael variants in proteins, and 94 variants in viral proteins.

*Acquiring Structural Models and Homology Models* We searched for crystal structures and homology models of proteins in **VTS** to maximize structural coverage. For proteins present in HumDiv without crystal structures in the PDB, we produced comparative models using Modeller(13, 14). For proteins with sufficient variant annotation details in UniProt but without structures in the PDB, we extracted comparative models from ModBase(39) and SwissModel(44), selecting models with the largest sequence identity match to the query. All protein models were standardized to remove unwanted components (duplicate chains, ligands, metals, and non-standard amino acids). This curation process resulted in 9,477 variants of 2,637 separate domains in 2,444 proteins (see Supplementary Figure S1).

### Protein Variant Characterization

Each protein variant is characterized by 106 features, five from sequence-based analysis, 17 from Rosetta ddg_monomer, 83 from Rosetta FastRelax, and one additional feature generated using PROBE.

*Sequence-based Features from BLAST Analysis* We find sequences similar to the query protein using PSIBLAST (2.2.25+, two iterations, pseudocount of two)(5) and extract five features directly from the output PSSM. At the protein position of interest, we use the PSSM log-likelihood of the native and variant amino acids (pssm_nat, pssm mut) along with the position’s information content (info_cont) as features. We also include an aminochange term that indicates broad chemical differences between the native and variant amino acid (see Supplementary Figure S2, Supplementary Table S4).

*Structure-based Features from Rosetta Analysis* Stability differences between the native and variant protein structures are predicted by comparing their individual Rosetta Energy terms(20). The Rosetta Energy function combines physical and statistical potentials to approximate the energetic stability of protein structures and can be decomposed into individual scoring terms(25). We derive structure-based features from two different approaches for refining the local structure around the new amino acid; a fast approach approximating the change in Energy (Rosetta ddg_monomer(20), **17** features) and broader conformational sampling using Rosetta FastRelax(21, 49) (**83** features). Both protocols 1) substitute the native residue for the variant amino acid, 2) refine the variant structure, including protein backbone movements, to accommodate this change, and 3) compare the output structures using the Rosetta score terms (Figure 1, Supplementary Figure S2). To generate features for each variant, we follow Poultney *et al.*(40) and normalize structure-based features by comparing scores for a given variant to scores derived from Rosetta-relaxed ensembles of its native protein. We also include the accessible surface area at the position of variation as a feature, calculated using PROBE(52). Additional details on the methods of structural analysis and generation of the 106 features can be found in the Supplementary Material.

**Figure 1.**
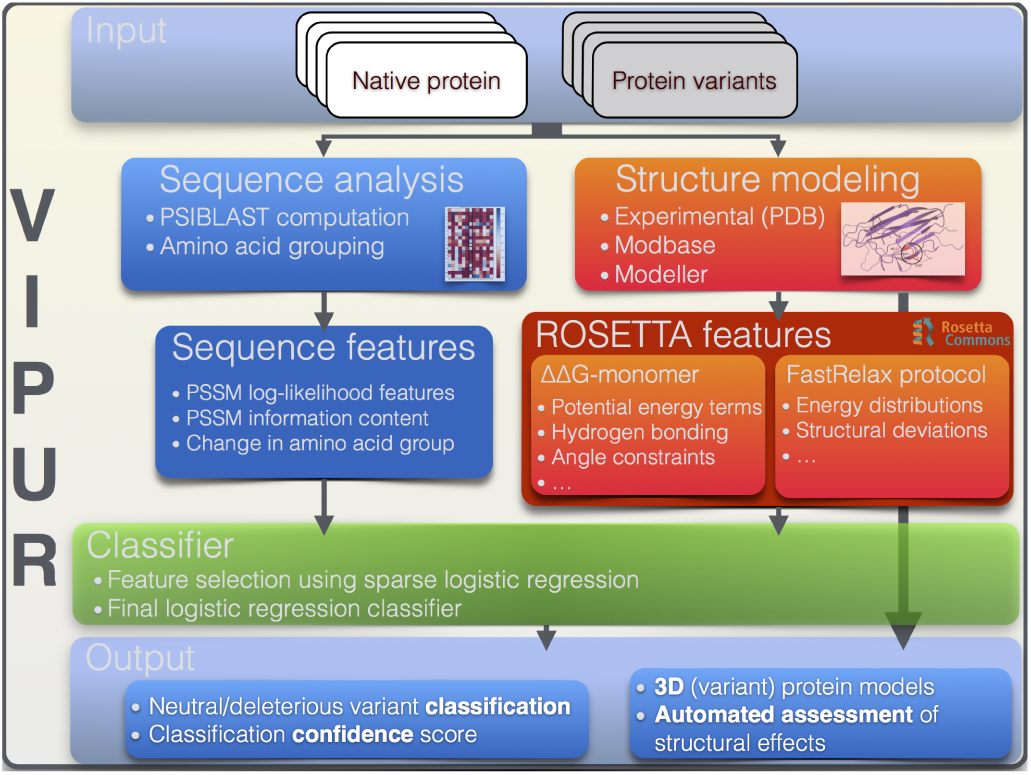
VIPUR Analysis Pipeline. Starting from a structural model of the native protein and a list of variants to be tested, VIPUR generates features using PSIBLAST and ROSETTA. Structure-based features are extracted from ROSETTA simulations comparing the native and variant protein structures. Variant structures are refined using the ddg_monomer protocol and the FastRelax protocol to consider a distribution of protein conformations. Features are combined in a logistic regression classifier that is trained on 9,477 variants from over 360 species. VIPUR outputs the predicted label (deleterious or neutral), a confidence score, the top scoring 3D models of the variant protein structure, and an automated interpretation of the variant effect derived from the weighted contributions of each feature to produce a physical description of protein disruption.

### Training a Sparse Logistic Regression Classifier

VIPUR uses sparse logistic regression as a statistical classification framework to robustly discriminate between **<u>deleterious</u>** and **<u>neutral</u>** protein variants from the derived 106 sequence-and structure-based features and thus allows for a natural probabilistic interpretation of the outcome. Using stability based feature selection we identified a set of 20 non-redundant features that maximize the average generalization performance(26, 31, 33) of the logistic regression classifier (Supplementary Table S4, Supplementary Figure S3, Supplementary Figure S4). We evaluated the performance of this classifier on 100 independent random splits (80% training, 20% testing, split by proteins not variants) by means of average Receiver Operating Characteristic and Precision-Recall curves (Figure 2, Figure S6). Using the same strategy we trained a sequence-only classifier using just the sequence-based features and a structure-only classifier using just the structure-based features (Supplementary Figure S4). We compared VIPUR curves to several alternative methods, including the individual sequence-based and structure-based feature sets (Supplementary Figure S5), an optimized SVM with a radial basis function kernel (Supplementary Figure S8), and PROVEAN (Figure 2). Many popular variant annotation methods are only benchmarked on human variants, making interpretation of their predictions non-applicable for nonhuman variants, such as **VTS**.

**Figure 2.**
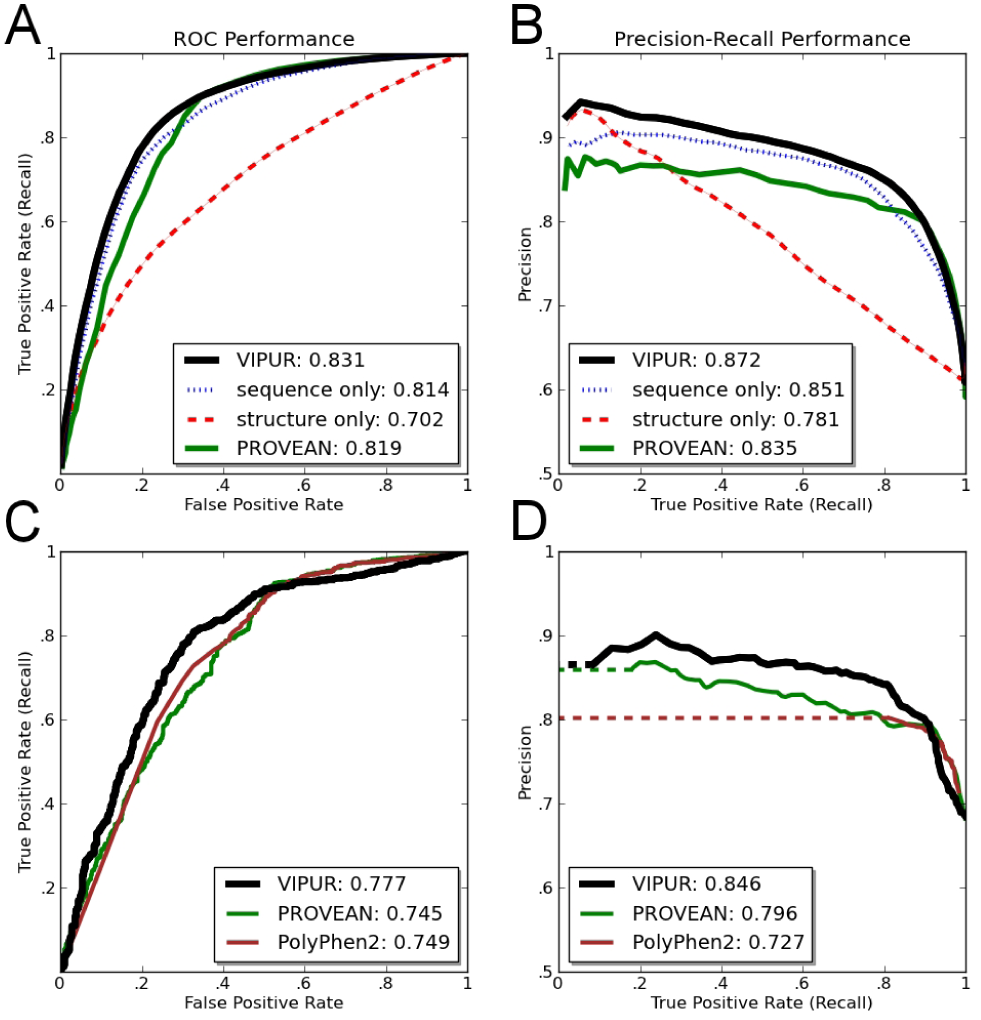
VIPUR Training ROC and PR Performance: Receiver Operating Characteristic and Precision-Recall curves for VIPUR and other popular methods. Curves (**A,B**) are averaged from 100 random splits (80% training, 20% testing) evaluated only on the leave-out testing sets. **A**) Our combined classifier (black) has increased specificity compared to PROVEAN (green) with comparable sensitivity and higher AUROC than all other methods tested. PROVEAN and our sequence-only classifier have very similar AUROC but appear to emphasize sensitivity and specificity respectively. **B**) VIPUR has notably increased AUPR over all other classifiers tested. Inclusion of the structure-based features improves classification (+2.5% accuracy) and dramatically improves ranking ability (.020 ΔAUPR). **C,D**) We cannot directly compare performance of VIPUR to classifiers trained on the same variants (HumDiv) or restricted to predictions of human proteins. A VIPUR-like classifier was trained using 7,935 variants from HumDiv and non-human proteins to compare performance with PolyPhen2 and PROVEAN on a set of 1,542 human variants. The VIPUR-like classifier achieves higher AUROC (**C**) and AUPR (**D**) than both PolyPhen2 and PROVEAN.

We cannot properly compare performance between VIPUR and PolyPhen2 on the full **VTS** since it contains variants in non-human proteins and variants from PolyPhen2’s training set (HumDiv). A set of 1,542 human variants included in **VTS** that are not included in HumDiv are used to compare a VIPUR-like classifier and PolyPhen2. To ensure a fair comparison, we retrained a VIPUR-like classifier (VIPUR^∗^) on the remaining 7,935 variants of **VTS**. We calculated ROC curves and PR curves for VIPUR^∗^, PolyPhen2, and PROVEAN on this set of 1,542 variants (Figure 2) and a subset of 383 variants found naturally in the human population (Supplementary Figure S7, Supplementary Table S3).

### VIPUR Software Availability

VIPUR is currently available as an independent Python module requiring BLAST+, ROSETTA, and PROBE (all freely available for academic use). Please see the VIPUR code for usage and analysis details, available at https://osf.io/bd2h4. The full predictions for all variants below, including structural models, are also available at <u>https://osf.io/bd2h4</u>.

### Classifying ClinVar Annotated Single Nucleotide Variants Phenotypes

We demonstrate that VIPUR’s **<u>deleterious</u>** predictions are an accurate indication of variant pathogenicity by classifying variants in the ClinVar database(24). ClinVar is a collection of human variants with annotated phenotypic effects, including variants with causative ‘**pathogenic’** effects and ‘**benign’** variants with no known disease effect. We expect VIPUR **<u>deleterious</u>** predictions to be enriched for variants with ClinVar “pathogenic”, “likely pathogenic”, or “risk factor” annotations, termed pathogenic variants. We also expect VIPUR **<u>neutral</u>** predictions to be enriched for variants with “benign” and “likely benign” annotations, termed **benign** variants. Many variants in ClinVar contain variants with **uncertain effects** or conflicting annotations (e.g. “likely benign” and “likely pathogenic”) including variants directly annotated with “uncertain significance”. We obtained models for 24,703 variants (in 4,016 proteins) in ClinVar from available structures in the PDB, ModBase, and SwissModel out of 32,311 variants (in 7,188 proteins) that could be unambiguously matched to UniProt proteins. ClinVar contains many additional SNV entries that lack appropriate protein IDs, variant positions, or annotations. Here we present predictions for VIPUR, PolyPhen2, and PROVEAN on 5,590 variants in ClinVar containing 498 benign variants, 1,797 pathogenic variants, and 3,295 variants with uncertain annotation. Additional predictions for CADD, SIFT, and PROVEAN were obtained from dbNSFP(28).

### Obtaining Inflammation Disease-Associated Variants

To demonstrate detailed VIPUR predictions on disease associated variants, we applied VIPUR to variants associated with inflammation diseases. We collected variants associated with various inflammation diseases and diabetes mellitus from entries in OMIM(30) and UniProt(2) by searching for the terms “Celiac disease”, “Crohn’s disease”, and “diabetes mellitus”, and mapping these variants onto available protein structures. This resulted in 388 variants in 46 disease-associated proteins. We provide illustrative examples of different **<u>deleterious</u>** variants and functional sites (Figures 4, 5, S9, S10).

### Classifying De novo Mutations in the Simons Simplex Collection

We tested VIPUR’s ability to identify disease-associated variants by classifying *de novo* missense mutations in the Simons Simplex Collection (SSC) of sequenced exomes from families (quads and trios) with children having Autism Spectrum Disorders (referred to as <u>probands</u>)(19, 37, 42) and unaffected <u>siblings</u>. Quad studies consist of exome sequencing for children with ASD, both of his or her parents, and siblings with no intellectual disability or ASD phenotype. These studies identify *de novo* SNVs in children with ASD (variants not present in either parent) and examples of *de novo* variation from the unaffected siblings. For 2,814 mutations in the SSC, 2,226 mutations could be analyzed by all variant annotation methods tested (1,335 missense mutations found in <u>proband</u> children and 891 mutations in their unaffected <u>siblings</u>). For VIPUR, 1,644 mutations were mapped onto structures from the PDB, ModBase, and SwissModel, considering models of all protein isoforms available for genes with alternative splicing. We predicted **<u>deleterious</u>** scores using VIPUR and applied our sequence-only classifier to the 582 mutations that could not be mapped to structure. For each mutation, we only considered the isoform prediction with the highest score, treating any **<u>deleterious</u>** prediction for a gene as indicative of deleteriousness. We compared the VIPUR, PolyPhen2 (HumDiv), and SIFT predictions to the phenotype associated with each *de novo* mutation (<u>proband</u> or unaffected <u>sibling</u>)(28). Since these methods have different scores, we consider the enrichment for proband mutations across score thresholds by calculating the ratio of proband to sibling mutations in different score bins. Although these classification methods differ, we expect high scores (**<u>deleterious</u>**, ‘damaging’, ‘intolerant’) to be enriched for <u>proband</u> mutations and low scores (**<u>neutral</u>**, ‘non-damaging’, ‘tolerant’) to be enriched for mutations found in unaffected <u>siblings</u> (Figure 6). We consider the correlation between this enrichment ratio and each output score across score thresholds and also the enrichment ratios found at the score cutoff of .5. Additional evaluation verified that this method of comparison is robust to the number of bins (Supplementary FigureS12) and the score threshold used (Supplementary FigureS13).

## RESULTS

### Accuracy and Generalization of the VIPUR Classifier

Combining sequence-based features and structure-based features enables VIPUR to accurately and precisely identify deleterious variants, achieving >90% precision on the highest scoring 40% (over 3,800 variants above score cutoff of .7, Figure 2B). VIPUR achieves a higher AUROC and AUPR than PROVEAN and other methods tested (Figure 2). Scores that clearly indicate confident predictions are essential for prioritizing variants and deleterious proteins. Filtering predictions with our confidence score raises the accuracy from 81% with no ranking (scores above .5 are considered **<u>deleterious</u>**) to > 94% accuracy for scores above .95. We tested both the classification (in-set) and generalization (out-of-set) performance of VIPUR and report here only the generalization performance (Figure 2) since this is characteristic of VIPUR’s behaviour on new variants. The classification and generalization performance converge as the training set size increases demonstrating that VIPUR predictions are robust and the classifier is not overfit to the training set (Supplementary Figure S4). Classifiers trained on only the sequence-based features correctly predict 78% of the dataset, providing a high baseline performance, while the structure-based features cause VIPUR output scores to scale with precision, indicating a clear estimate of prediction confidence. Adding structure-based features improves performance by recovering improperly classified **<u>neutral</u>** samples with a slight change in **<u>deleterious</u>** sensitivity, suggesting these features help identify *misclassifications* made by the sequence-based features (Figure 2A).

### Comparison to Other Classifiers

We compare performance of our combined classifier to PROVEAN, PolyPhen2, and multiple classifiers trained on our own features (structure and sequence features only). PROVEAN is a popular variant annotation method that extends the SIFT framework for identifying deleterious variants. We compare performance on the entire **VTS** to PROVEAN since it can interpret variants in any organism without additional training and is not overfit to any particular training set. Using the full **VTS** our combined classifier performs better than PROVEAN with improved classification (AUROC 0.831 over PROVEAN’s 0.819) and notably improved ranking ability, quantified by our AUPR of 0.872 over PROVEAN’s 0.835; over twenty percent of the AUPR not covered by PROVEAN (Figure 2A). Our sequence-only classifier displays similar performance to PROVEAN, with nearly identical AUROC (Figure 2). The “flat” shape of the precision-recall curves for sequence-based classifiers may be a general property of these feature sets, providing generalized predictions without clear specificity since they do not identify any specific mechanism of protein disruption. These similarities also suggest that our sequence-based features appropriately capture the deleterious signal within multiple sequence alignments (when used with logistic regression).

We are unable to consistently compare performance of VIPUR to popular human-specific methods on the full **VTS**. For example, PolyPhen2 does not support prediction on nonhuman variants and is trained on HumDiv (contained in **VTS**). Accordingly, we compare our method to PolyPhen2 over a subset of 1,542 human variants in **VTS** using a classifier similar to VIPUR but trained on the remaining 7,935 variants of **VTS**, termed VIPUR^∗^. VIPUR^∗^ produces ROC curves similar to PROVEAN and PolyPhen2 with notably improved AUPR on this set of human variants (Figure 2C,D). PROVEAN and PolyPhen2 perform very similarly although PolyPhen2 predictions are restricted to a small region of the Precision-Recall landscape (PolyPhen2 scores are highly degenerate, a large number of predictions obtain a score of ‘1’). The decrease in performance for VIPUR and PROVEAN on this set of variants suggests these variants represent mutations that are different from the rest of **VTS**. VIPUR^∗^ appears overfit, due to the lack of diverse neutral annotations during training (HumDiv neutrals are all pseudomutations) and we included all available variants with neutral annotations to eliminate this overfitting when training VIPUR. We also contrast the performance of our logistic regression classifier with a SVM classifier using an optimized Radial Basis-Function kernel (Supplementary Figure S6). Our logistic regression classifier achieves higher accuracy, AUPR, and AUROC than the SVM classifier with fewer features (reduced complexity), superior generalization, and direct interpretability (Supplementary Figure S6B).

We investigated prediction trends of VIPUR across numerous protein properties including the source of data, species of origin, fold classification, functional annotation, and model quality (using Pearson chi-squared test, see Supplementary Material). These trends show a slightly increased false negative rate for eukaryotic proteins and a slightly increased false positive rate for prokaryotic proteins. This is likely caused by simple label imbalance since the majority of neutral-labeled variations are in eukaryotic proteins. While VIPUR generalizes very well across diverse protein functions, the structure-only classifier has an increased false negative error rate on several DNA and RNA associated proteins, suggesting that simulating these interactions will improve the accuracy of our structural modeling (DNA and RNA are absent in our structural models). We have verified that VIPUR’s performance is the same for proteins with many variants in **VTS** and proteins with no other variants in the training set. This demonstrates that VIPUR is not overfit to specific sequence/fold properties, a confounding form of overfitting(16) (Supplementary Table S2).

### VIPUR Predictions Match ClinVar Phenotypes

We tested VIPUR’s capability to distinguish **pathogenic** variants from **benign** variants by classifying SNVs in the ClinVar database. ClinVar’s curated annotations include **benign** variants with no known effect on disease and a large collection of **pathogenic** variants with various causal roles in genetic disorders and disease susceptibility. The variant annotations in ClinVar do not directly match VIPUR labels, but we expect ClinVar **pathogenic** variants to be enriched for **<u>deleterious</u>** VIPUR predictions and for ClinVar **benign** variants to be enriched for **<u>neutral</u>** VIPUR predictions. We emphasize that *not all* pathogenic variants are deleterious and many deleterious variants appear benign when they do not have clear biological phenotypes.

**Pathogenic** variants have a highly skewed distribution of VIPUR **<u>deleterious</u>** scores while **benign** variants have a broad distribution of **<u>neutral</u>** scores (Figure 3). PolyPhen2 scores tend towards high and low values that also clearly distinguish between **pathogenic** and **benign** variants. PROVEAN scores are distributed similarly to VIPUR scores, matching our expectations for ClinVar variants. All three methods are designed to highlight deleterious variants and must be able to clearly identify variants with strong evidence of deleteriousness. Benchmarked on ClinVar, VIPUR has a higher specificity than PolyPhen2 with a reduced sensitivity. ClinVar itself has a high label bias with a 7:2 proportion of **pathogenic:benign** annotations. Training on datasets with a large label imbalance can inherently off-set the sensitivity/specificity tradeoff of a classifier and must be avoided by training on samples that *accurately* represent the category labels. Prediction methods of other variant annotation methods resemble VIPUR predictions and match our expectations for score distributions on **pathogenic** and **benign** variants (Supplementary Figure S11).

**Figure 3.**
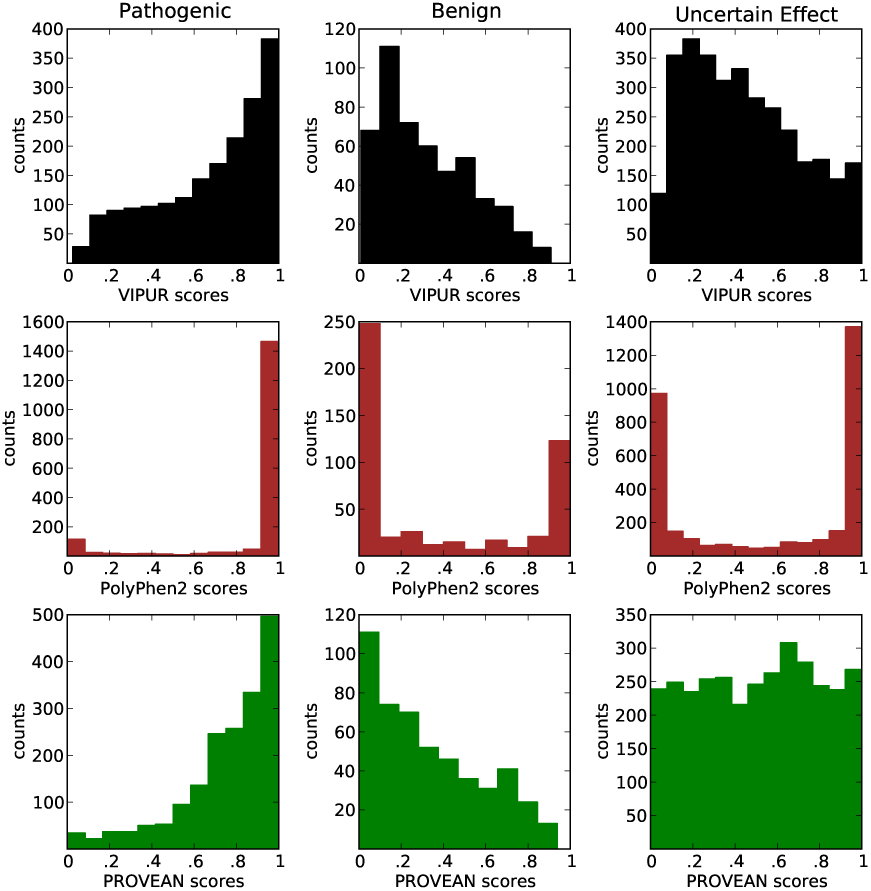
VIPUR Scores Clearly Identify Pathogenic Variants: VIPUR predictions on ClinVar variants match expectations from their phenotype annotations. left) **Pathogenic** variants have a skewed distribution of VIPUR **<u>deleterious</u>** scores (>.5) and are correctly predicted by PolyPhen2 and PROVEAN. center) **Benign** variants have a broad distribution of VIPUR **<u>neutral</u>** scores (<.5) while PolyPhen2 pushes variants to high and low scores. In contrast to the high **pathogenic** label bias of ClinVar, we expect most genetic variations to be **benign** and unlikely to disrupt protein function. right) Predictions on ClinVar variants annotated with **uncertain effect** highlights VIPUR’s ability to identify a small set of likely **<u>deleterious</u>** variants while PolyPhen2’s high false positive rate leads to an overwhelming number of high confidence “probably damaging” predictions. VIPUR’s score distribution resembles the **benign** variants with a small set of confident **<u>deleterious</u>** predictions while PROVEAN scores are uniformally distributed.

Predictions on ClinVar variants annotated as **uncertain effect** demonstrate the differences in error rates between these methods (Figure 3, Supplementary Figure S7). VIPUR predictions are predominately **<u>neutral</u>** with a small set (208/3,295, 6%) of highly confident **<u>deleterious</u>** predictions while PolyPhen2 predicts over seven times as many “high confidence” pathogenic variants (1,435/3,295, 44%!). PROVEAN predictions are nearly uniform without enrichment at the highest and lowest scores or a score distribution resembling either **benign** or **pathogenic** variants. Without reliable labels for ClinVar variants of **uncertain effect**, the accuracy of these predictions cannot be evaluated. VIPUR is the only method tested with a score distribution for these variants resembling the **benign** variants and places the fewest number of these variants into the highest confidence bins (Supplementary Figure S11, Supplementary Table S5). Nearly all of these methods identify some aspect of deleteriousness although classification of variants with **uncertain** labels is very diverse between these methods. Several variant annotation methods may have artificially high false positive rates and comparisons between these methods will obtain similar score distributions when benchmarked on datasets with a large deleterious label bias (like ClinVar). The **uncertain effect** variants likely have a different label ratio, leading to the diverse behavior of these methods.

### Examples of Detailed Structural Annotation for Deleterious Variants Associated with Human Diseases

We demonstrate VIPUR’s applications by predicting **<u>deleterious</u>** variants among a small set of inflammation and diabetes associated variants. Genome Wise Association Studies and exome sequence studies of disease conditions reveal many candidate genes by associating variants to traits and conditions. Some of these genes with be **<u>deleterious</u>** and may have large effects on the disease phenotype. The variants collected here do not necessarily have causal roles in inflammation or diabetes, unlike the ClinVar **pathogenic** variants which have established effects, but instead provide examples of VIPUR prioritization and interpretation. We collected proteins and variants associated with the terms “Celiac disease”, “Crohn’s disease”, and “diabetes mellitus” from OMIM(30) and UniProt(2), identifying 388 variants in 46 disease-associated proteins (in 102 models). We predicted VIPUR scores for each variant and interpreted the structure-based features of each variant model. Predictions on the entire set of disease-associated variants are available at <u>https://osf.io/bd2h4</u>.

Out of 388 variants, we predict 205 are **<u>deleterious</u>** with 108 having confidence scores above .8. UniProt annotations for these deleterious variants have several keywords describing damaging effects. These descriptions, however, do not meet our curation standard for a **<u>deleterious</u>** label in VTS but are suggestive of the variant’s functional impact. Our physically intuitive structure-based features allow VIPUR to automatically produce structural hypotheses about the physical causes of deleteriousness. We include a summary of the structure-based features that contribute to the deleterious classification with each prediction, indicating disrupted hydrogen bonds, disulfide bridges, improper packing, and other structural defects. Many deleterious variants destabilize the protein native state by introducing a steric clash or otherwise preventing proper packing arrangements. In this dataset, variants in NR3C1, HNF1A, NEUROD1, and SIAE all clearly disrupt packing interactions. During classification, features like the Rosetta van der Waals repulsive term (fa_rep) contribute a large deleterious score, allowing automated identification of packing disruption. While these amino acid changes dramatically alter the side-chain shape and size, amino acid side-chain interactions are most easily identified using 3D contacts in the protein structure. VIPUR’s structure-based features automatically detect disrupted side-chain interactions using Rosetta’s statistical potentials. In this dataset, variants in LEP, AKT2, and TGM2 are predicted to disrupt specific interactions that stabilize the folded protein. These examples are representative of automated VIPUR interpretations but many long-range effects require sampling protein backbone conformations to properly interpret variant effects.

Many physical interactions within a protein are far apart in sequence, limiting the insight provided by methods that assume protein positions are independent. VIPUR can correctly identify mutations that disrupt these interactions by analyzing a 3D structural model of the protein, even when destabilization occurs far from the mutated position. We identified several cases where mutations disrupted interactions between elements of secondary structure, a deleterious effect captured by VIPUR but missed by sequence-based methods. The S204P variant of IL6 is associated with numerous inflammation diseases (Figure 4) and annotated in UniProt as “87% loss of activity”. While PROVEAN predicts this variant is neutral (-1.20 score), VIPUR predicts this variant is **<u>deleterious</u>** with high confidence (.835) and infers that it disrupts a disulfide bond. Position 204 is not close enough to destabilize the nearest disulfide bond, C101-C111, by direct interaction (Figure 4, bottom), however, conformational rearrangements that accommodate P204 disrupt the interface between helix four and helix seven, straining this disulfide bond. These subtle structural changes cannot be detected with a multiple-sequence alignment or structural modeling of a single conformation. V117M of ADIPOQ also appears neutral in a PSSM and PROVEAN (-2.00 score), but interactions between protein backbones with *β*-strand pairing inform a **<u>deleterious</u>** prediction by VIPUR (Supplementary Figure S9). V117 is physically close to I135 on an adjacent *β*-strand and mutation of V117 to methionine introduces a clash between these positions that cannot be accommodated without breaking inter-strand hydrogen bonds, destabilizing the *β*-sheet (Supplementary Figure S9, bottom right). These examples demonstrate the clarity and scope of structural modeling to detect destabilizing mutations, highlighting the limited performance of sequence-based methods at positions without strong conservation.

**Figure 4.**
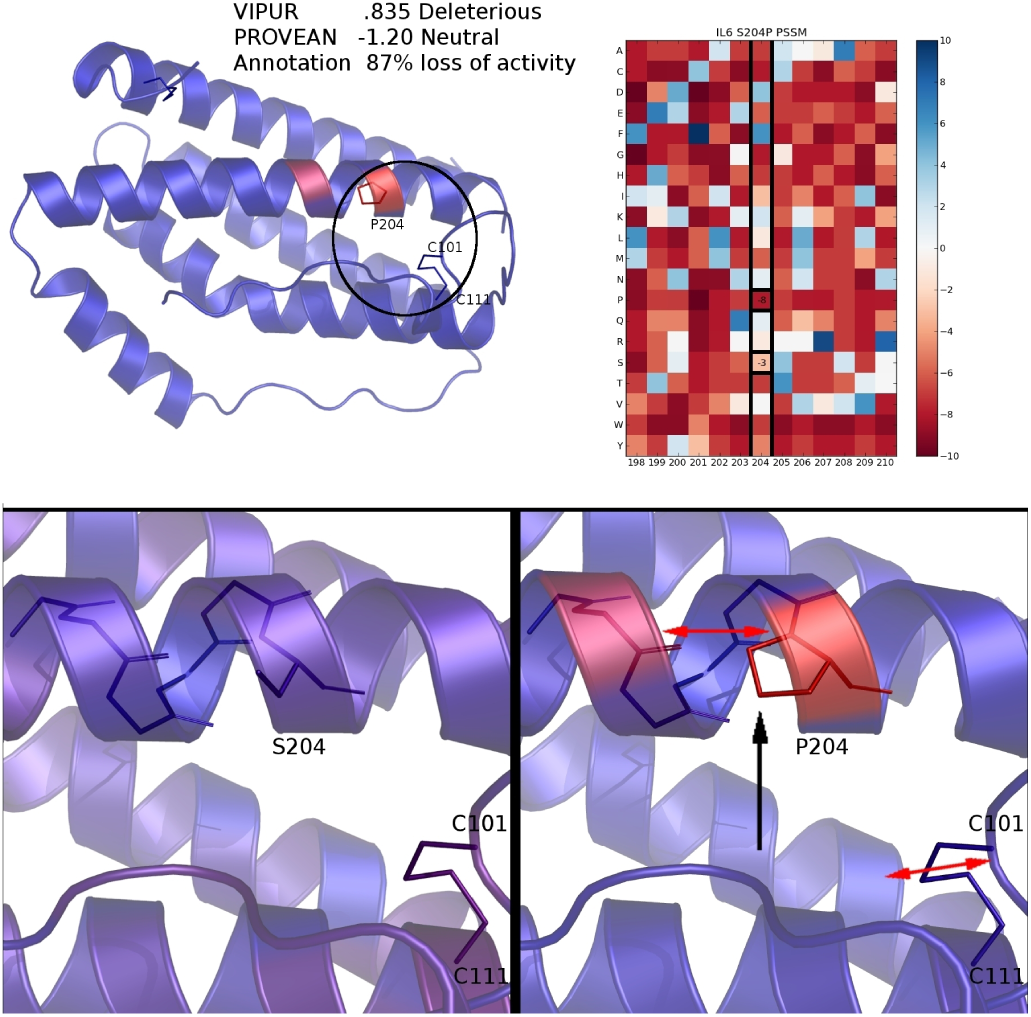
S204P disrupts a critical helix interface in IL6. VIPUR predicts S204P is **<u>deleterious</u>** (.835), matching the UniProt annotation “87% loss of function” and infers the **<u>deleterious</u>** label due to destabilized disulfide bond, while PROVEAN predicts S204P is neutral (-1.20 score). Every residue in IL6 is colored by the difference in Rosetta energy between the native and variant protein structures, highlighting the destabilization introduced by S204P (top left). The PSSM generated by PSIBLAST does not indicate strong conservation for serine at position 204 (top right, PSSM columns shown for surrounding residues). The native S204 structure has a stable interface (bottom left, residues colored by Rosetta energy of a representative model) but becomes destabilized in the P204 variant model (bottom right). Perturbing this helix could accommodate the proline destabilization, however, this strains the nearby C101-C111 disulfide bond (bottom right), leading to an accurate **<u>deleterious</u>** prediction.

Beyond long-range interactions, VIPUR can also detect destabilization at active sites and binding interfaces. GCK has many diabetes-associated variants, including several high confidence predictions in this dataset: T168P, G299R, W257R, and G385V. Position 168 is a conserved glycine in the PSSM and predicts both the native threonine and variant proline are similarly unfavorable. This conservation causes PROVEAN to predict T168P as deleterious (-5.82) even when the native threonine is just as disfavorable as the variant (based on sequence analysis), yet known to make a hydrogen bond with the substrate D-glucose (from PDB 3F9M). Our structural model does not include this interaction with D-glucose (all ligands are removed) yet VIPUR still predicts mutation to proline is highly destabilizing (.987) based on the unfavorable backbone conformation of proline (Figure 5) at this *structurally conserved* binding site. We observe a similar pattern at other ligand and metal binding sites, such as ZFP57 H374D (not shown), where structure-based features produce confident **<u>deleterious</u>** predictions even without explicitly including the ligand or metal in the structural model. This suggests interaction sites have conserved structural properties that can help identify deleterious variants and that VIPUR predictions may identify disrupted active sites even when the substrate is unknown.

**Figure 5.**
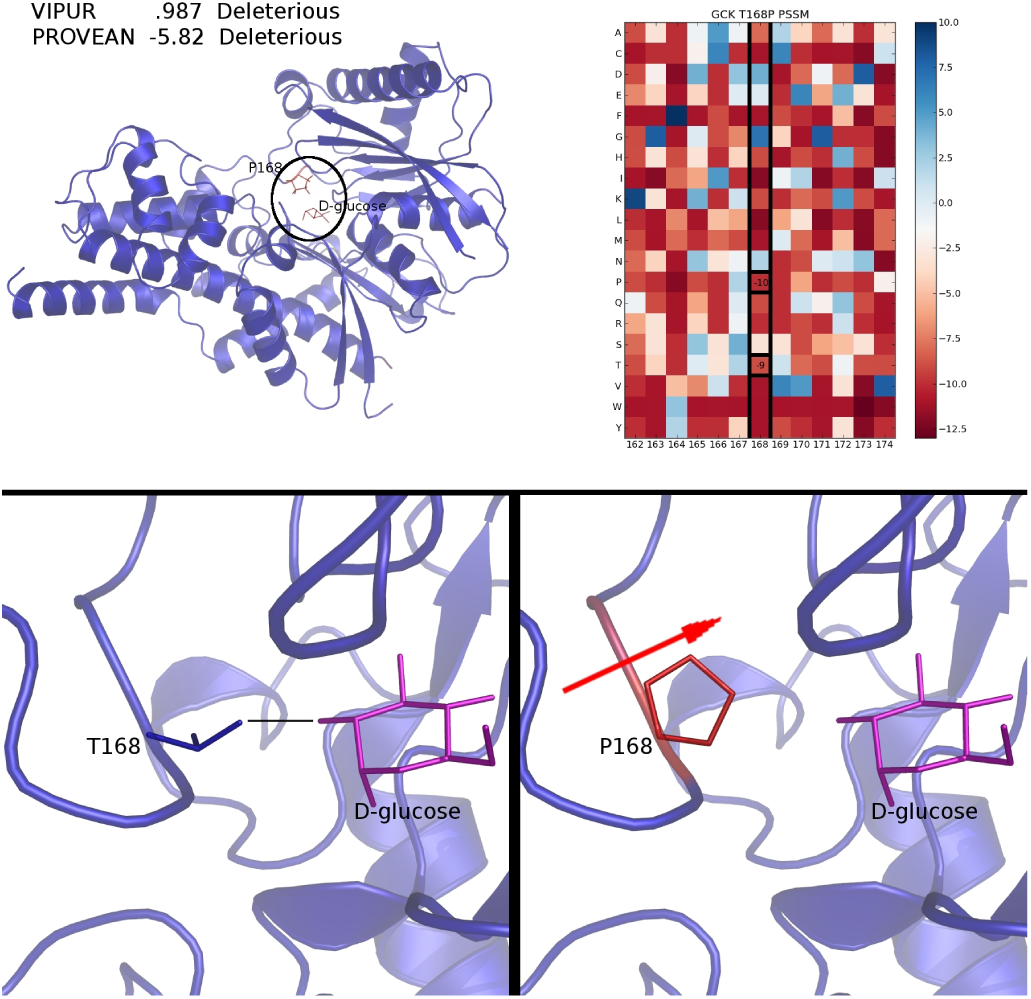
T168P destabilizes an active site loop in GCK. T168P is predicted **<u>deleterious</u>** (.987) due to disrupted backbone interaction (hydrogen-bonding) and is also predicted **<u>deleterious</u>** confidently by PROVEAN (-5.82). Every residue in GCK is colored by the difference in Rosetta energy between the native and variant protein structures, highlighting the high energy of P168 (top left). The PSSM generated by PSIBLAST indicates both the native and variant amino acids are not favored at position 168 (top right, PSSM columns shown for surrounding residues). The native T168 forms a hydrogen bond to the substrate, D-glucose (bottom left, ligand position from PDB 3F9M, residues colored by Rosetta energy of a representative model), which is absent in the P168 variant. Although interaction with D-glucose is not simulated during classification, VIPUR predicts proline is destabilizing due to disrupted backbone hydrogen bonding, suggesting other active sites can be accurately classified even without bound ligands.

### Identification of Deleterious De novo Mutations Associated with Autism Spectrum Disorders

To demonstrate VIPUR’s ability to prioritize disease-associated genetic variants in the absence of curated labels, we ran VIPUR on the Simons Simplex Collection.

The Simons Simplex Collection (SSC) is a set of *de novo* SNVs where the genotypes of children with Autism Spectrum Disorders are compared to their parents, identifying *de novo* variation. These quad studies require genomic comparison to both parents, the child with ASD, and an unaffected sibling to provide samples of *de novo* variation found in children without ASD. Many of the variants in the SSC may be *non-causal* for ASD or otherwise contribute weak effects to complex behavioural phenotypes, obscuring the deleteriousness and pathogenicity of these variants. We expect the **<u>deleterious</u>**/damaging/intolerant predictions from these methods to be enriched for *de novo* mutations found in children with ASD (probands) while **<u>neutral</u>**/no effect/tolerant predictions are enriched for variants in unaffected <u>siblings</u>.

**Table 1.** Predictions on the Simons Simplex Collection. proband-D: proband mutations in deleterious predictions (True Positives) at .5 cutoff, proband-N: proband mutations in neutral predictions (False Negatives) at .5 cutoff, sibling-D: sibling mutations in deleterious predictions (False Positives) at .5 cutoff, sibling-N: sibling mutations in neutral predictions (True Negatives) at .5 cutoff, proband enrich: the ratio of proband-D/sibling-D, sibling enrich: the ratio of proband-N/sibling-N

| 2,226 de novo variants from the SSC (1,335 proband, 891 sibling, 1.50 label bias) |  |  |  |  |  |  |
| --- | --- | --- | --- | --- | --- | --- |
| method | BLOSUM62 | VIPUR | SIFT | PolyPhen2 | CADD | MutationTaster |
| proband-D | 907 | 554 | 779 | 769 | 747 | 830 |
| proband-N | 428 | 781 | 556 | 566 | 588 | 505 |
| sibling-D | 566 | 348 | 499 | 494 | 499 | 546 |
| sibling-N | 325 | 543 | 392 | 397 | 392 | 345 |
| # > .95 | 0 | 43 | 545 | 607 | 186 | 1340 |
| proband enrich | 1.60 | 1.59 | 1.56 | 1.56 | 1.50 | 1.52 |
| sibling enrich | 1.32 | 1.44 | 1.42 | 1.43 | 1.50 | 1.46 |
| Spearman | 0.90 | 0.87 | 0.54 | 0.53 | 0.13 | 0.07 |
| Spearman p-value | 0.08 | **2.68e-3** | 0.11 | 0.12 | 0.73 | 0.91 |
| Pearson | 0.92 | 0.86 | 0.44 | 0.49 | 0.03 | 0.11 |
| Pearson p-value | **0.03** | **1.39e-3** | 0.20 | 0.15 | 0.93 | 0.81 |

These methods all output confidence scores that are scaled from 0 to 1 with high scores predicting deleterious effects and low scores predicting neutral effects. When thresholding prediction scores at .5, all methods tested have a higher proportion of proband mutations in deleterious predictions and a lower proportion in neutral predictions, however none of the methods appear notably enriched. Since the classification threshold is arbitrary, no single threshold will be appropriate for all methods, however, we expect <u>proband</u> enrichment to be proportional to the confidence score. We count the number of <u>proband</u> and <u>sibling</u> mutations found in each score bin and compare this ratio to the confidence score of that bin. We calculate the correlation between the annotation confidence score and <u>proband</u> enrichment to compare method performance.

The simple BLOSUM62 matrix achieves an impressive enrichment for <u>proband</u> mutations despite having only seven distinct values for mutations in this dataset. Surprisingly, PolyPhen2, SIFT, CADD, and MutationTaster do not display significant enrichment across score thresholds, although SIFT and PolyPhen2 have trends in the proper direction for intolerant/damaging predictions (Figure 6). VIPUR is the only method to obtain significant Spearman (rank) and Pearson correlations across score thresholds, properly enriching **<u>deleterious</u>** predictions for <u>proband</u> mutations and removing <u>proband</u> mutations from **<u>neutral</u>** predictions. VIPUR predictions also fit our intuition that the majority of variants in this dataset are predicted to have a **<u>neutral</u>** effect on protein function. Many of these variant effect annotation methods are trained and/or benchmarked on datasets with a high label bias. This label imbalance likely contributes to the inflated false positive rate we observe for many methods tested here (Supplementary Figure S7). Since we are primarily concerned with the efficient identification of candidates for follow-up studies, proper *ranking* of pathogenic variants is essential for highlighting causal mutations and is severely confounded by these high false positive rates for *de novo* mutations. At the confidence score threshold of .95, VIPUR predicts 43 variants are very likely to have disrupted molecular functions which may contribute to ASD while PolyPhen2 predicts 607 variants with high confidence. While these confidence thresholds are arbitrary, we verified that this trend is invariant to the number of bins used (Supplementary Figure S12) or the classification thresholds used (Supplementary Figure S13).

**Figure 6.**
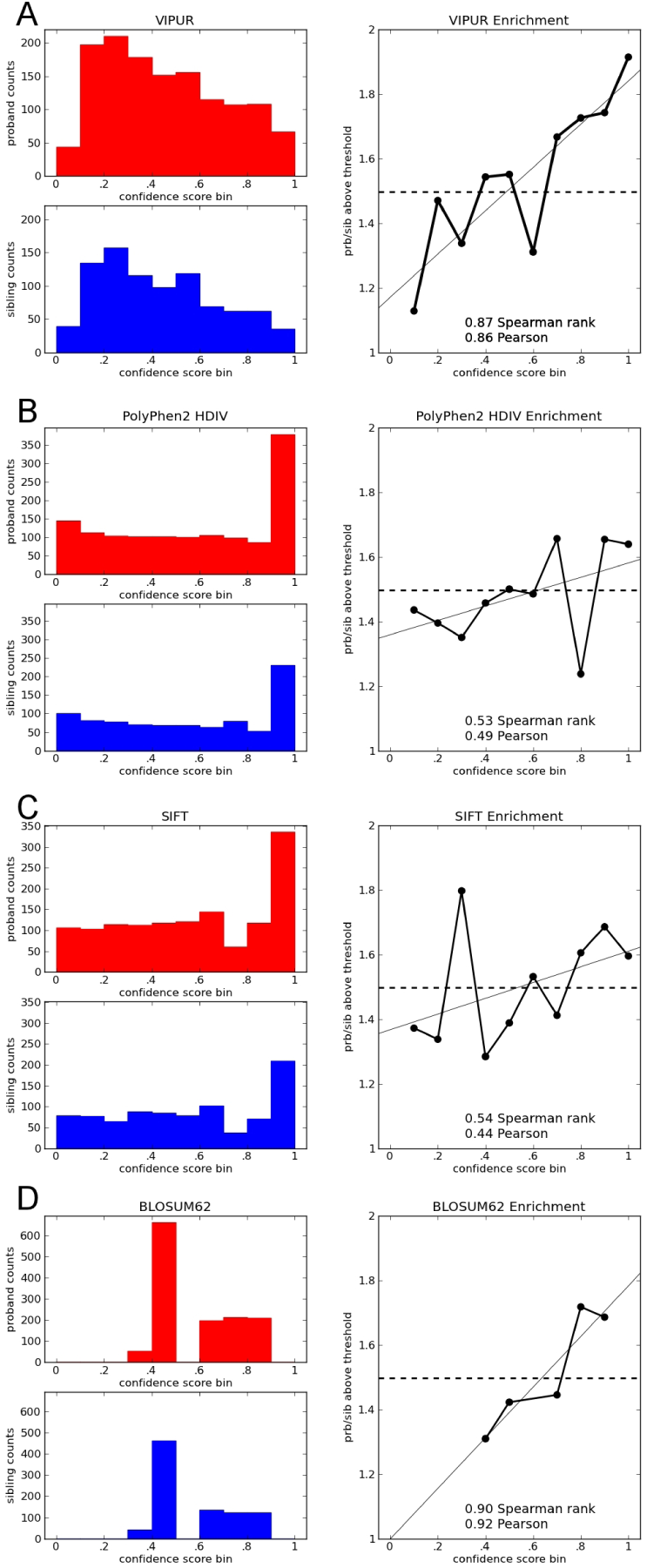
VIPUR Deleterious Predictions Identify Autism-associated Mutations: Predictions for various methods on the Simons Simplex Collection, containing 1335 de novo mutations found in children with autism-spectrum disorders (probands) and 891 de novo mutations found in unaffected siblings. For each prediction method, the distribution of confidence scores is shown for proband (red) and sibling (blue) mutations. The ratio of these counts for each score bin are shown along with the background expectation (dashed line, 1.50, 1335/891). We expect high deleterious scores to be enriched for proband mutations and low scores enriched for mutations found in siblings. A) VIPUR predicts most mutations in both sets have neutral effects and properly enriches for proband mutations at high scores *and* de-enriches for probands mutations at low scores. B) PolyPhen2 effectively splits mutations into a high confidence bin vs everything else, however this top bin is not strongly enriched for proband mutations. C) SIFT scores are distributed similarly to PolyPhen2 with similar overall correlation, however its fluctuation around the background expectation are different. D) Using the simple BLOSUM62 score (negative scores are deleterious) yields an excellent enrichment for proband mutations, however the scores are not truly continuous leading to fewer scores (smaller p-value).

## DISCUSSION

VIPUR is a variant annotation method that is designed to identify deleterious variants by analyzing conservation and protein structural energetics. The VIPUR **<u>deleterious</u>** and **<u>neutral</u>** labels are learned from curated annotations of variants with clear effects on protein molecular functions and are not restricted to variants with known pathogenicity for any particular disease or any single organism. VIPUR has superior performance to PROVEAN and PolyPhen2 on out-of-set evaluations drawn from **VTS**. Our structure-based features enhance the ranking ability of VIPUR, leading to an improved precision for variants with higher **<u>deleterious</u>** scores. We demonstrate that VIPUR predicted labels match expectations for the **pathogenic** and **benign** phenotype annotations in the ClinVar database. All other variant annotation methods tested also match these expectations as well, although the methods notably disagree about ClinVar variants with **uncertain effect**. Examples of VIPUR predictions on inflammation and diabetes associated variants demonstrate the clarity of structure-based features to explain the specific causes of protein deleteriousness. These automated structural interpretations are only possible using structure refinement techniques that can identify long-range structural disruption. Predictions on the Simons Simplex Collection show that VIPUR **<u>deleterious</u>** predictions are more enriched for *de novo* mutations found in children with autism spectrum disorders than any other method tested. While **VTS** and ClinVar have a strong deleterious label bias, we expect most genetic variations to have neutral effects and VIPUR consistently predicts neutral scores for collections of variants of unknown significance.

Our current method allows us to accurately predict and interpret many protein variants, however several substantial improvements to this method are on the horizon. Successful prediction of variants in IL6 (Figure 4) and ADIPOQ (Figure S9) demonstrate that VIPUR can accurately predict the effects of amino acid substitutions even when disruption occurs at a distant region of the protein structure. This suggests that VIPUR could predict the functional effects of multiple mutations within the same protein, even though these variants are not currently included in **VTS**. Thus far **VTS** includes 323 of the 400 possible single amino acid transitions. Although we observe nearly unbiased predictions across these amino acid transitions, VIPUR has slightly reduced performance for some substitutions with changes in polarity (Supplementary Table S1). More advanced electrostatics modeling in the context of our predicted structure ensembles will likely improve classification for these transitions(18). In addition to more sophisticated electrostatic features, many additional features are likely to improve performance, such as individual amino acid properties. Recent improvements to the Rosetta framework make it possible to incorporate DNA, RNA, metals, and other cofactors into our structural models which will further improve our structure-based features and interpretation. Improved Rosetta protocols for modeling membrane environments, including transmembrane-specific conformational sampling and a membrane energy function with depth dependent solvation and hydrogen bonding terms, will expand our coverage to include variants in transmembrane environments(3, 53).

**VTS** currently includes 9,477 annotated variants in more than 360 species with 106 features for each variant and structural models from the Protein Data Bank and homology models. Independent of VIPUR, this dataset is a valuable resource for researchers in computational biology and machine learning communities to develop and test novel classification methods. We are currently expanding **VTS** to include annotated variants with multiple substitutions, nearly neutral variations, variants in transmembrane proteins(3), alternative comparative models using multi-template homology modeling(13), and known binding interactions including variants at DNA-and RNA-protein interfaces. These advances will make VIPUR applicable to an even wider range of protein variants, further contributing to our understanding of structure-function relationships. Given the relatively distinct chemical environments and conformational motions between intrinsically disordered protein regions, transmembrane proteins, and traditional ordered proteins, we expect individual classifiers trained for each type of protein region will perform better than a marginal classifier trained on all types combined. While the PDB does not include models of all proteins, human proteins are abundant and available models in ModBase and SwissModel help increase the structural coverage. Of the 32,311 protein coding variants in ClinVar (in 7,188 proteins) that could be unambiguously matched to proteins in UniProt, 24,703 (in 4,016 proteins) had structures available in the PDB, ModBase, or SwissModel (76% of variants covered, 55% of proteins). We apply our sequence-only classifier to protein variants lacking structural models and will continue to improve this rapid classification method. Although structural coverage limits our ability to classify all protein variants, VIPUR still identifies candidate genes and causal variants within large genomic datasets, highlighting only the variants with structural evidence of large effects.

## CONCLUSION

VIPUR has been designed to identify and interpret deleterious protein variants across multiple species and sources of variation. To achieve this generalization, we have collected and curated **VTS**, a dataset of protein variants with annotated functional and physical effects on protein molecules. VIPUR’s superior classification performance and ranking stem from a seamless integration of high quality sequence and structure information (Figure 2) and Rosetta’s ability to find low energy backbone conformations that can accommodate neutral substitutions and indicate long-range disruption of deleterious substitutions. Unlike other methods, VIPUR uses automated structural analysis to make a detailed 3D model of each variant and subsequently infer the physical origin of deleterious predictions, generating hypotheses and interpretations previously achievable only by tedious manual inference. We have demonstrated that VIPUR predictions are informed by protein structural constraints that cannot be identified using a multiple sequence alignment or a static protein structure alone (Figure 4, Figure 5). VIPUR can automatically highlight protein variants involved in human diseases that disrupt protein function and is applicable to nonsynonymous SNVs in proteins with reliable structural models. Although VIPUR predicts variants with disruption of biophysical function, this label matches expectations of biological phenotypes and predicts fewer false positives than many current variant annotation methods, a problem confounded by the incoherence of label bias between traditional benchmarks (more pathogenic examples than neutral examples) and real applications (we expect most single variants to be neutral). While other methods lack the specificity required to identify neutral variation, VIPUR can clearly distinguish deleterious variants from neutral variants (Figure 6A, B). Previous advances in deleterious variant prediction have often focused on improving recall and global accuracy but failed to explain the origin of deleterious variation. Here, we demonstrate how these pathogenicity detection methods are great tools for initially filtering and identifying *potential* causal variants, however additional analysis, such as structural model analysis, is required to further refine candidates. VIPUR can identify deleterious protein variants and provide structural explanations for disrupted protein function. We hope that VIPUR will contribute to our understanding of structure-function relationships, particularly for the interpretation of de novo mutations and disease associated variants.

## ACKNOWLEDGEMENTS

We would like to thank the Simons Foundation, specifically the Simons Foundation Autism Research Initiative and the Simons Center for Data Analysis, and NYU-ITS, specifically Muataz Al-Barwani and the NYU Abu Dhabi ITS. RB was supported by the Simons Foundation and US National Science Foundation grants IOS-1126971, CBET-1067596 and CHE1151554, and National Institutes of Health GM 32877-21/22, PN2-EY016586, IU54CA143907-01 and EY016586-06.*Conflict of interest statement.* None declared.

## SUPPLEMENTARY MATERIAL

### VTS Acquisition and Curation

*Label Curation* We collected protein variants (nonsynonymous SNPs) from HumDiv(1) and UniProt(2) with clear deleterious or neutral effects. These variants were mapped onto crystallographic and comparative models of the protein macromolecules from the Protein Data Bank(4), ModBase(39), and SwissModel(44). HumDiv is a database of naturally occurring human protein variants annotated as causing Mendelian diseases and used for calibration and testing numerous prediction tools(1). We extracted additional variant annotations from UniProt(2) using the curation rules of HumDiv. Variants with annotations describing clear evidence that some molecular activity essential to the protein function is disrupted are labeled ‘**deleterious’** (if the activity is reported, only activity ≤ 5% is labeled deleterious). Variants with annotations describing clear evidence that all known molecular activities essential to the protein function are unperturbed are labeled ‘**neutral’** (if the activity is reported, only activity ≥ 70% is labeled neutral). If there is insufficient evidence, we do not assign either label, including annotations with clear effects that do not guarantee disruption of molecular function (ex. no annotation, disease-associated, lethality, low expression, improper localization, etc.).

*Acquiring Structural Models and Homology Models* We searched for crystal structures and comparative models of proteins in the dataset to maximize coverage. For proteins present in HumDiv without crystal structures in the PDB, we produced comparative models using Modeller(13, 14). We restricted our templates to structures generated using X-ray crystallography with more than 20% sequence identity to the query protein, selecting templates with the highest sequence identity match to the query. When a single template could not cover all variant positions (e.g. missing densities), multiple models were constructed (e.g. separate domains) or multiple templates were used to cover these missing regions. Comparative models were produced using Modeller for threading(14), skipping refinement steps that would be redundant with refinement during feature generation. Many of these comparative models had obvious structural defects, such as broken loops and improbable backbone Φ-Ψ angles, requiring curation of over 5,000 putative models. For proteins with sufficient variant annotation in UniProt but without structures in the PDB, we extracted comparative models from ModBase(39) and SwissModel(44) (no restriction on the template sequence identity to query), selecting models with the largest sequence identity match to the query. All protein models were standardized to remove unwanted coordinates (duplicate chains, ligands, metals, and nonstandard amino acids). We removed all structures covering transmembrane regions since our current Rosetta analysis does not appropriately sample or score transmembrane regions. This curation process resulted in 9,477 variants in 2,637 models of 2,444 proteins (see Supplementary Figure S1).

*Structure-based Features From Rosetta Analysis* The ddg_monomer protocol is designed to approximate the change in free energy upon mutation (ΔΔG) and uses a fast refinement protocol which outputs the change in Rosetta Energy (stability)(20), contributing 17 features to our analysis. Rosetta FastRelax uses Monte Carlo sampling of protein backbone conformations with side chain optimization(49) to find low Energy conformations. We run Rosetta FastRelax to generate 50 low Energy conformations(20) for both the native and variant proteins. We include additional features describing the geometric differences between the input and final structure for each trajectory (e.g., RMSD and gdtmm) to detect proteins undergoing large rearrangements, totaling 23 features. To compare the native and variant ensembles and eliminate potential differences in score magnitude across diverse protein folds, we 1) extract the distributions of each Rosetta score term for the native and variant proteins, 2) calculate the quartiles of the variant protein score distributions, and 3) calculate the cumulative density for these quantiles on the corresponding native protein score distribution(40). FastRelax and quartile analysis produce three features per score term for each variant, corresponding to the Q1, Q2, and Q3 quartiles(40), totaling 60 features.

*VIPUR Software Implementation and Availability* VIPUR is currently available as an independent Python module and requires BLAST+, ROSETTA, and PROBE. VIPUR runs on a structural model of the native protein structure (in PDB format) and a file containing the variants to predict (e.g. S204P), or an entire directory of these files. VIPUR verifies the positions and native amino acids of the variant file and controls execution of PSIBLAST(5), Rosetta ddg_monomer(20), Rosetta FastRelax(49), and PROBE(52). The output includes predictions from the VIPUR classifier and classifiers using only structure-based features and sequence-based features, and an interpretation of the variant effect including the top ranking structural features. Please see the VIPUR code for full usage and analysis details, available at <u>https://osf.io/bd2h4</u>.

### Assessment of VIPUR Classifier Training and Performance

*Feature Selection Using Sparse Logistic Regression* VIPUR uses logistic regression as a statistical classification framework to robustly discriminate between **deleterious** and **neutral** protein variants from the derived 106 sequence-and structure-based features. Logistic regression generalizes linear regression to binary classification by linking known class labels (*y_i_*, **deleterious** vs. **neutral**) to our feature set (vectors x*_i_*) using a logistic function and allows for a natural probabilistic interpretation of the classification outcome (Prob(*y_i_*|x*_i_*)). Consider our dataset by 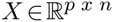 where *p*=106 is the number of features and *n*=9,477 denotes the number of labeled variants. For each column (variant) characterized by features x*_i_* ∈*X* we have a binary class label *y_i_* ∈[−1,1] where the positive label indicates a deleterious variation. These class labels, derived from curated annotations, are stored in the vector 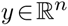. We learn

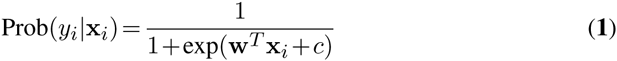

where Prob(*y_i_*|x*_i_*) estimates the conditional probability of label *y_i_* given the sample x*_i_*. The model is characterized by a weight vector 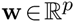 and bias/intercept 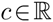. This model provides separation by learning the w*^T^*x*_i_* + *c* = 0 hyperplane in the feature space, where Prob(*y_i_*|x*_i_*)=0.5. Thus, w*^T^*x*_i_* + *c* > 0 corresponds to a **deleterious** prediction with individual weight *w_j_* corresponding to the relative importance of the feature for making this separation. We determine w by finding the minimum of the associated negative log-likelihood (also called the logistic loss) on the training data 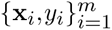.

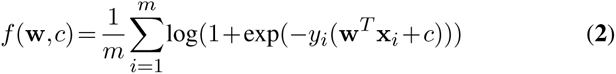

To arrive at a classifier with a minimal, non-redundant feature set that simultaneously generalizes well across all protein variants, we repeatedly split the dataset into training and test sets (100 random splits) and compute logistic regression models across all model complexities (i.e., using one to 106 features). We use sparsity-promoting L1-regularization(26, 33) to reduce complexity, seeking the minimum of this loss function, *f*(w, *c*) while simultaneously promoting sparsity of the weight vector w as described below:

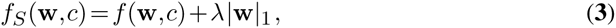

where |·|_1_ denotes the L1 norm and *λ* > 0 is a parameter promoting sparsity (tunable). This sparsity implies that only a few features are used to predict the class labels (most weights *w_j_* “shrink” to 0). For any fixed *λ* the convex non-smooth loss *f*_S_(w, *c*) can be efficiently solved using projected sub-gradient methods.

The sparsity constraint restricts the number of features selected during training, but does not ensure features are robustly selected. Similar to stability selection in linear regression (31), we recorded the frequency of features that are present in the best predictive model on each test set. We determined the model complexity (number of features) that maximizes the average generalization performance to be 20and thus select the 20most frequent features (Figure 2B, Supplementary section). Samples were split by proteins, rather than variants, to prevent the classifier from learning protein-specific patterns(1). For each split, we tuned the sparsity constraint (parameter *λ*) and selected the model that minimized error on the testing set. We generated and tested 100 of these splits and recorded the features selected for every model (Supplementary Figure S4). Features were ranked according to their prevalence in the trained models of these splits, selecting 20 features that maximize generalization performance (Figure 2, Supplementary Figure S4) and perform better than models trained on all 106 features. We tested generalization performance for classifiers trained on increasing fractions of the full dataset to determine robustness. Classification and generalization performance approach very similar values when trained on only 50% of the data, demonstrating models learned from these 20 are generalizable and not overfit (Figure 2B, Supplementary Figure S4). The final VIPUR classifier was trained on the full dataset using only these 20 selected features. We evaluated the performance of the final sparse logistic regression classifier on 100 independent random splits (80% training, 20% testing) by means of average Precision-Recall and Receiver Operating Characteristic curves (Figure 2, Figure S6). VIPUR performs better than several alternative methods, including an optimized SVM with a radial basis function kernel (Supplementary Figure S8).

*VIPUR Performance Breakdown and Biases* To ensure our classifier was trained on a sufficiently diverse set of protein variants, we integrated naturally occurring variation with variants produced by mutagenesis and pseudomutations derived from differences between humans and closely related mammals(1). There is an abundance of protein-specific data available for the proteins in our dataset, however, we restricted our features to information/analyses available for under-researched proteins (hence benchmarking with comparative model structures). We investigated prediction trends of VIPUR for numerous protein properties including the source of data, species of origin, structural context, functional annotation, and model quality to identify any biases in our predictions and suggest which sources of information may improve VIPUR further. For each of these protein “subsets”, we tested if performance on these variants had biased accuracy, error rates, or composition. We considered groups based on: data source (HumDiv or UniProt), model source, domains, species, structural context (surface, core etc.), GO molecular function, GO biological process, and amino acid transitions using a Pearson chi-squared test, restricting our inquiry to groups with at least 100 samples in the data. Here, we comment on the most significant deviations, correcting for cases where skewed predictions occurred on imbalanced samples (ex. surface variants have a higher number of neutral predictions and also have a higher number of **neutral** samples).

**Supplementary Table S1.**
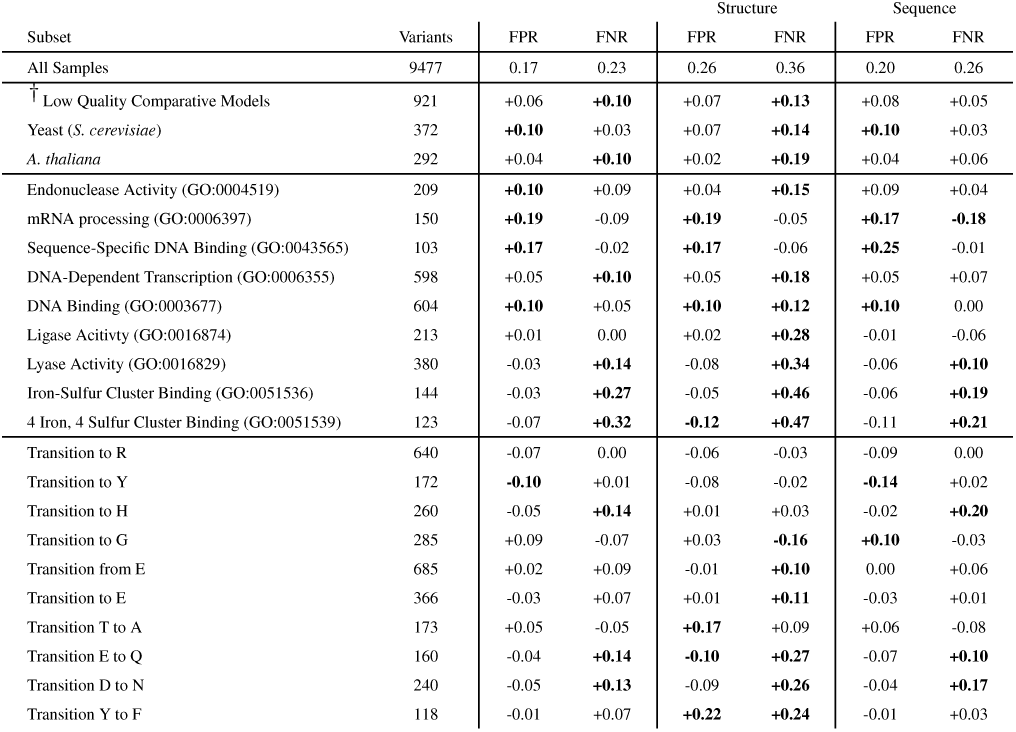
Performance Trends. FPR = False Positive Rate, FNR = False Negative Rate, ^†^comparative models from templates with <30% sequence identity to the query Error Rates more than .10 above expectation are in **bold**

There is no identifiable difference in performance between variants from HumDiv and UniProt despite HumDiv containing solely human data. Surprisingly, we also detect no difference in performance between models drawn from the PDB, ModBase, SwissModel, or produced by Modeller (not shown). As expected, lower quality comparative models perform worse than high quality models (models from templates under 30% sequence identity are reduced to 73% accuracy). These lower quality models have higher false negative rate, suggesting that low quality models do not capture the interactions near the variant position that are destabilized upon mutation. We do not observe a difference in performance between crystal structures and high quality comparative models, although model defects may already be masked by our structure-based features which always consider the differences between native and variant models. Our dataset contains examples for 323 of the 400 possible canonical amino acid transitions and we detect a slightly higher false negative rate for transitions to histidine and transitions from glutamic acid to lysine.

Across organismal domains, eukaryotic proteins have a slightly increased false negative error rate while prokaryotic proteins have a slightly increased false positive rate. This trend may be caused by label imbalance as the majority of **neutral**-labeled variations are eukaryotic. Human proteins are predicted more accurately that other species, likely due to label imbalance since most **neutral** annotations are in human proteins. Proteins from yeast (*S. cerevisiae*) and *A. thaliana* have slightly higher error rates than expected. We do not detect any difference in performance for variants on the surface or interior of proteins, although labels and predictions are both imbalanced with more **neutral** examples on the protein surface and more **deleterious** examples in the protein core.

We observe a higher than expected false positive rate among proteins with DNA binding functions including sequence-specific DNA binding (GO:0043565), DNA-dependent transcription (GO:0006355), and cell cycle (GO:0007049) annotations. Although we expected the structure-based features to perform worse for proteins associated with nucleic acids (due to the absence of these molecules in the structural model), this functional bias comes from both feature sets. A similar bias occurs for transitions to glycine and may represent a tendency to classify all variations at highly conserved sites as **deleterious**, independent of the new amino acid’s properties. We do not observe biases for other nucleic acid functions (nucleotide binding, RNA binding, nuclease activity) however, we see a higher false positive rate for endonuclease activity (GO:0004519) and mRNA processing (GO:0006397) annotations in both feature sets. The structure-based features have higher false negative rates for catalytic activity (GO:0003824), specifically ligase (GO:0016874) and lyase (GO:0016829) activity annotations, however, all of these trends are reduced in the combined classifier.

We anticipated that many active sites in our structural models would lack chemical interactions necessary for correct classification since ligands and cofactors are not included. However, several of these positions have highly constrained conformations that allow accurate **deleterious** predictions with structure-based features even without the ligand present (Figure 5). We also observe an increased false negative rate for iron-sulfur associated proteins (GO:0051536, GO:0051539), and a slight decrease in performance for metal binding proteins (GO:0046872). As with ligands, this decrease in performance was much less than anticipated (Supplementary Table S1), likely caused by the highly constrained geometries of metal coordination sites that are captured by our structure-based features, even without the metal(s) present. We observe several amino-acid specific biases, mostly for the structure-only classifier, indicating areas for improvement. Our classifier predicts mutation to arginine and to tyrosine correctly for nearly all examples available. The structure-based features have a higher false positive rate for the transitions: glutamic acid to glutamine, aspartic acid to asparagine, threonine to alanine, and tyrosine to phenylalanine. These transitions may involve subtle water coordination sites, and generally indicate that performance could be improved with more rigorous electrostatics methods.

Nearly all of the trends for sequence-based features involve higher false positive rates while trends for the structure-based features involve higher false negative rates. Since mutations at conserved positions are more likely to be **deleterious** than non-conserved positions, sequence-based analysis is more likely to falsely label variation at conserved sites as **deleterious** (too sensitive), independent of the new amino acid’s properties. Since our structural models do not include binding partners (proteins, DNA, ligands, metals), this structure-based analysis is missing some interactions and limited to predicting **deleterious** variants that cause energetic disruption of the monomer. When combined, these two sources of information provide a clear interpretation of conservation and destabilization, allowing the combined confidence metric to scale directly with performance and correctly classify variants missed by both feature sets independently. Many of these trends suggest areas for improvement, particularly the use inclusion of nucleic acid models and more sophisticated electrostatic methods for polar amino acid characterization.

*VIPUR Avoids Confounding Sources of Circular Predictions* A recent study of pathogenicity prediction methods showed that several classifiers trained on mutational data made circular predictions, effectively learning a majority vote rule for specific protein families(16). This circularity highlighted the difficulty to learn generalized classifiers on data with a high label bias (e.g. when there are many more **deleterious** samples than **neutral** ones) and suggested methods for avoiding and testing for circularity. Unlike other similar classifiers, VIPUR was trained on a highly curated set of variants with experimentally validated labels. This curation process eliminated many variants with ambiguous effects, limiting the number of proteins that contribute multiple mutations and reducing the label bias to ≈3/5 (5,740/3,737).

To prevent any circularity from inflating evaluation of VIPUR, we ensured that all divisions of training and testing sets were striated by the protein identity, such that all variants from any individual protein are contained only in a single training or testing set (e.g. never training and testing on variants from the same protein). This guarantees that our performance metrics are evaluated on variants in proteins that had never been seen before. Assessment of circularity can be done by investigating any performance differences between variants in proteins with only a single variant in the training set and those with multiple(16). **VTS** contains 1125 ‘single’ variants in proteins that contain no other variants in the dataset, 3475 ‘pure’ variants in proteins with multiple variants that all share the same label, and 4877 ‘impure’ variants in proteins with multiple variants containing at least one from both labels (**deleterious** and **neutral**). Both the ‘single’ and ‘pure’ categories are enriched for **deleterious** labels while the ‘impure’ variants are enriched for **neutral** labels, suggesting an interesting bias in the data; that **neutral** samples are rarely provided without accompanying **deleterious** samples. Confounding circularity occurs when training on pure variants leads to overestimation on single and impure variants, creating a classifier that does not properly generalize. VIPUR performance on each of these variant groups is nearly identical to the others and the overall assessment (Supplementary Table S2). As the proportion of **deleterious** and **neutral** variants narrows for ‘impure’ proteins, there is a notable drop in performance metrics, particularly AUPR, however this likely stems from the high label bias in these subsets (mostly neutral samples). We also compared these performance trends to a simple per protein Majority Vote classifier where variants were simply assigned the majority label of other variants in the same protein (‘pure’ and ‘single’ variant predictions are uninteresting). This classifier has a similar trend of decreasing performance as the stricter label ratios are imposed, however even the best majority vote prediction has a similar performance to the worst performing subset of VIPUR predictions. These methods of assessing circularity are useful but may overpenalize any supervised learning classifier. VIPUR predictions have similar performance trends for variants with many identical labels in the dataset and variants in proteins that have never been seen before, avoiding overfitting by circularity.

**Supplementary Table S2.**
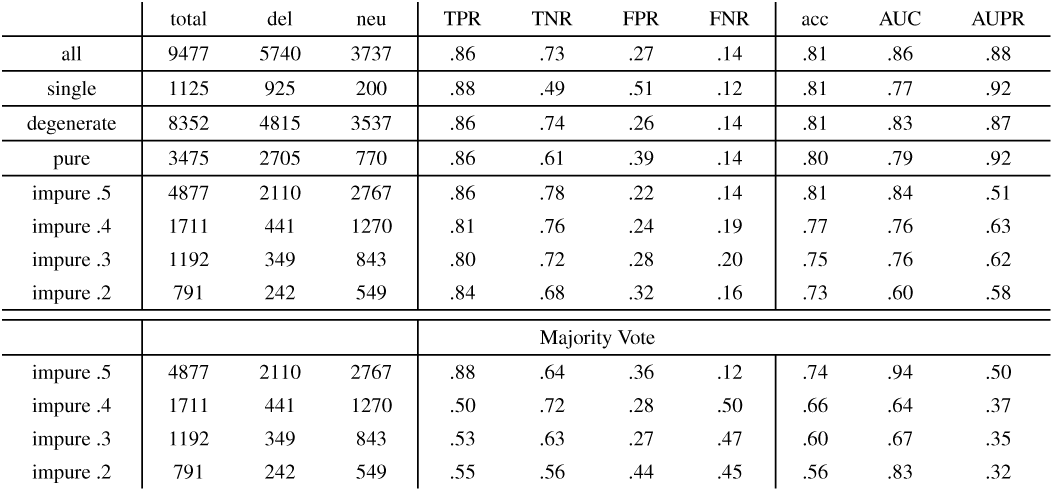
VIPUR Avoids Circularity. single: variants in proteins that contain no other variants, degenerate: variants in proteins with at least one other variant, pure: variants in proteins with at least one other variant with all variants sharing the same label, impute X: variants in proteins with at least one other variant and a label ratio of .5 ± X

|  | total | del | neu | TPR | TNR | FPR | FNR | acc | AUC | AUPR |
| --- | --- | --- | --- | --- | --- | --- | --- | --- | --- | --- |
| all | 9477 | 5740 | 3737 | .86 | .73 | .27 | .14 | .81 | .86 | .88 |
| single | 1125 | 925 | 200 | .88 | .49 | .51 | .12 | .81 | .77 | .92 |
| degenerate | 8352 | 4815 | 3537 | .86 | .74 | .26 | .14 | .81 | .83 | .87 |
| pure | 3475 | 2705 | 770 | .86 | .61 | .39 | .14 | .80 | .79 | .92 |
| impure .5 | 4877 | 2110 | 2767 | .86 | .78 | .22 | .14 | .81 | .84 | .51 |
| impure .4 | 1711 | 441 | 1270 | .81 | .76 | .24 | .19 | .77 | .76 | .63 |
| impure .3 | 1192 | 349 | 843 | .80 | .72 | .28 | .20 | .75 | .76 | .62 |
| impure .2 | 791 | 242 | 549 | .84 | .68 | .32 | .16 | .73 | .60 | .58 |
| Majority Vote |  |  |  |  |  |  |  |  |  |  |
| impure .5 | 4877 | 2110 | 2767 | .88 | .64 | .36 | .12 | .74 | .94 | .50 |
| impure .4 | 1711 | 441 | 1270 | .50 | .72 | .28 | .50 | .66 | .64 | .37 |
| impure .3 | 1192 | 349 | 843 | .53 | .63 | .27 | .47 | .60 | .67 | .35 |
| impure .2 | 791 | 242 | 549 | .55 | .56 | .44 | .45 | .56 | .83 | .32 |

### Comparison to Other Prediction Methods

*Comparison to PolyPhen2* PolyPhen2 and many popular pathogenicity detection methods are restricted to classification of human protein variants. We designed VIPUR to run on variants found in any species and avoid overfitting any particular organism. Eliminating this overfitting is particularly important when classifying *de novo* mutations since these variants are likely to be absent from multiple sequence alignments, reducing the accuracy of sequence-based analysis. We cannot properly compare performance between VIPUR and PolyPhen2 on the full **VTS** since it contains variants in non-human proteins and variants from PolyPhen2’s training set (HumDiv). There are 1,542 human variants in **VTS** that are not included in HumDiv and to ensure a fair comparison with PolyPhen2, we retrained a VIPUR classifier (VIPUR^∗^) on the remaining 7,935 variants of **VTS**. We calculated ROC curves and PR curves for VIPUR^∗^, PolyPhen2, and PROVEAN on this set of 1,542 variants and a subset of 383 variants found naturally in the human population (383 variants).

**Supplementary Table S3.** Prediction Performance on Human Variants in the VTS. TP = True Positives, FN = False Negatives, FP = False Positives, TN = True Negatives, Acc = Accuracy, Bal Acc = Balanced Accuracy (equal weight for both label classes), AUROC = Area Under the Receiver-Operating Curve, AUPR = Area Under the Precision Recall curve

| 1,542 variants (1,051 deleterious, 491 neutral, 2.14 label bias) |  |  |  |  |  |  |  |  |
| --- | --- | --- | --- | --- | --- | --- | --- | --- |
|  | TP | FN | FP | TN | Acc | Bal Acc | AUROC | AUPR |
| VIPUR* | 888 | 166 | 199 | 289 | 0.762 | **0.716** | **0.777** | **0.846** |
| PROVEAN | 973 | 78 | 265 | 226 | **0.778** | 0.693 | 0.745 | 0.796 |
| PolyPhen2 | 992 | 59 | 306 | 185 | 0.763 | 0.66 | 0.749 | 0.727 |
| 383 variants (311 deleterious, 72 neutral, 4.32 label bias) |  |  |  |  |  |  |  |  |
| VIPUR* | 249 | 65 | 22 | 47 | 0.767 | 0.730 | 0.797 | **0.929** |
| PROVEAN | 279 | 32 | 27 | 45 | 0.846 | **0.761** | **0.810** | 0.914 |
| PolyPhen2 | 291 | 20 | 34 | 38 | **0.859** | 0.732 | 0.805 | 0.858 |

VIPUR^∗^ produces ROC curves similar to PROVEAN and PolyPhen2 with notably higher specificity and a slightly reduced sensitivity (ROC and PR curves cross, Supplementary Figure S7). PROVEAN and PolyPhen2 perform very similarly although PolyPhen2 predictions are restricted to a small region of the Precision-Recall landscape since many predictions obtain scores of ‘0’ or ‘1’ (Supplementary Figure S7B,C). While this limitation of PolyPhen2 scores restricts the inferences on Precision-Recall performance, this reflects a practical limitation when identifying causal variants from GWAS and exome studies caused by degeneracy of the output metric. Across 1,542 human variants, VIPUR^∗^ has higher AUROC (Supplementary Figure S7A) compared to PROVEAN and PolyPhen2 with a notably higher AUPR (Supplementary Figure S7B). These variants have a notable label bias and are drawn primarily from mutagenesis of human proteins, resembling *de novo* mutations. When performance is evaluated on a subset of 383 variants (with notably higher label bias) that are found within the human population, PROVEAN and PolyPhen2 performance increases notably, although VIPUR^∗^’s ranking ability remains superior. These 1,542 variants are included in the training set of VIPUR to ensure it generalizes to mutagens and to eliminate artificially inflated performance metrics.

*Obtaining PROVEAN Predictions and Ranking* We compared performance of our predictive model to PROVEAN, a highly accurate method for labeling protein variants as “**damaging**” or “**neutral**”. For human variants, PROVEAN was run remotely using the (PROVEAN server at <u>http://provean.jcvi.org/index.php</u>). For non-human variants, PROVEAN was run locally using CD-hit 4.5.4. The output PROVEAN score does not have a clear increase in accuracy for high scoring predictions. We scaled PROVEAN scores between 0 and 1 over the range of scores obtained on this dataset with 0 being the smallest score and 1 being the largest score. Precision-recall and ROC curves are generated from *ranked* predictions so this scaling should have no impact on performance.

*Training and Optimizing a Support Vector Machine Classifier* We compared the performance of our logistic regression classifier to an optimized support vector machine using the entire feature set. Training and testing was performed using LibSVM (version 3.1) through Weka (3.6.0). We used a radial basis function kernel optimized over C ∈{2^−1^, 2^0^, 2^1^, &,2^9^} and *γ* ∈{2^−8^, 2^−7^, &,2^−1^}. Optimal parameters (C=2^2^, *γ* = 2^−4^) were selected using 5-fold cross-validation.

We evaluated performance using AUPR and AUROC for 100 splits of 80% training and 20% testing. Probability estimates for each prediction are determined in LIBSVM using a form of Platt et al.(17).

### Comparison of Prediction Methods on De novo Mutations Associated with Autism Spectrum Disorders

We evaluated VIPUR predictions for 2,815 *de novo* mutations in the Simons Simples Collection. Predictions for PolyPhen2, CADD, SIFT, and MutationTaster were obtained from dbNSFP(28). To ensure prediction performance is comparable, we reduced evaluation to 2,226 variants that have predictions available for all these methods. We calculated the enrichment of **deleterious** predictions for proband mutations by comparing the ratio of proband mutations to mutations in unaffected siblings across different **deleterious** score cutoffs. We expect high confidence **deleterious** predictions to be more enriched for proband mutations and low confidence **deleterious** predictions to be enriched for mutations from unaffected siblings. This performance trend is evaluated by calculating the correlation between each method’s deleteriousness scores and the enrichment for proband mutations. Proband or sibling association does not establish deleteriousness, however we expect proband mutations to have more **deleterious** mutations producing a large positive correlation for these prediction methods. The correlation values reported in the main text are calculated using 10 bins and this parameter will alter the exact correlation values obtained, but not the overall trend (Figure S12).

Independent of the number of bins used for calculating the correlation, VIPUR predictions correlate more highly with proband enrichment than other methods (Supplementary Figure S12). Since the number of bins is arbitrary, smaller bin sizes produce increasingly undersampled correlation estimates (fewer samples each bin) with most methods having a notable drop after 10 bins. Several methods do not notably change behaviour between 3 and 10 bins so we have reported Spearman rank correlation and Pearson correlation values using 10 bins. Independent of the threshold used to define **deleterious** and **neutral** classifications, VIPUR provides higher enrichment than other methods for **deleterious** predictions and lower or comparable enrichment for **neutral** predictions (Supplementary Figure S13). Most methods fluctuate around the background enrichment ratio, but VIPUR, PolyPhen2, and SIFT all have the expected enrichment trends, although VIPUR notably outperforms the other methods. Deleterious prediction proband enrichment using VIPUR sharply increases above .5 and this threshold is used as the default boundary for **deleterious-neutral** definition to fairly represent VIPUR’s enrichment. PolyPhen2 and SIFT follow very similar prediction patterns for these robustness evaluations which may suggest a deeper correlation between these methods.

Correlation values were tested for significance using the R software package. The reported p-values are compared to the null hypothesis that each correlation is zero. While BLOSUM62 values provide a very accurate baseline, they are not continuously distributed and can only be separated into 7 different score bins, reducing the overall significance of this correlation. CADD produces several metrics and has no recommended confidence cutoff for identifying deleterious mutations so we used the scaled Raw Rank Score (performs better than CADD Raw Score on this data). We compared predictions on *de novo* mutations to PolyPhen2 HumDiv (rather than PolyPhen2 HumVar) since it is designed to classify rare alleles and shares training data with VIPUR.

## Additional Methods Details

**Supplementary Table S4.**
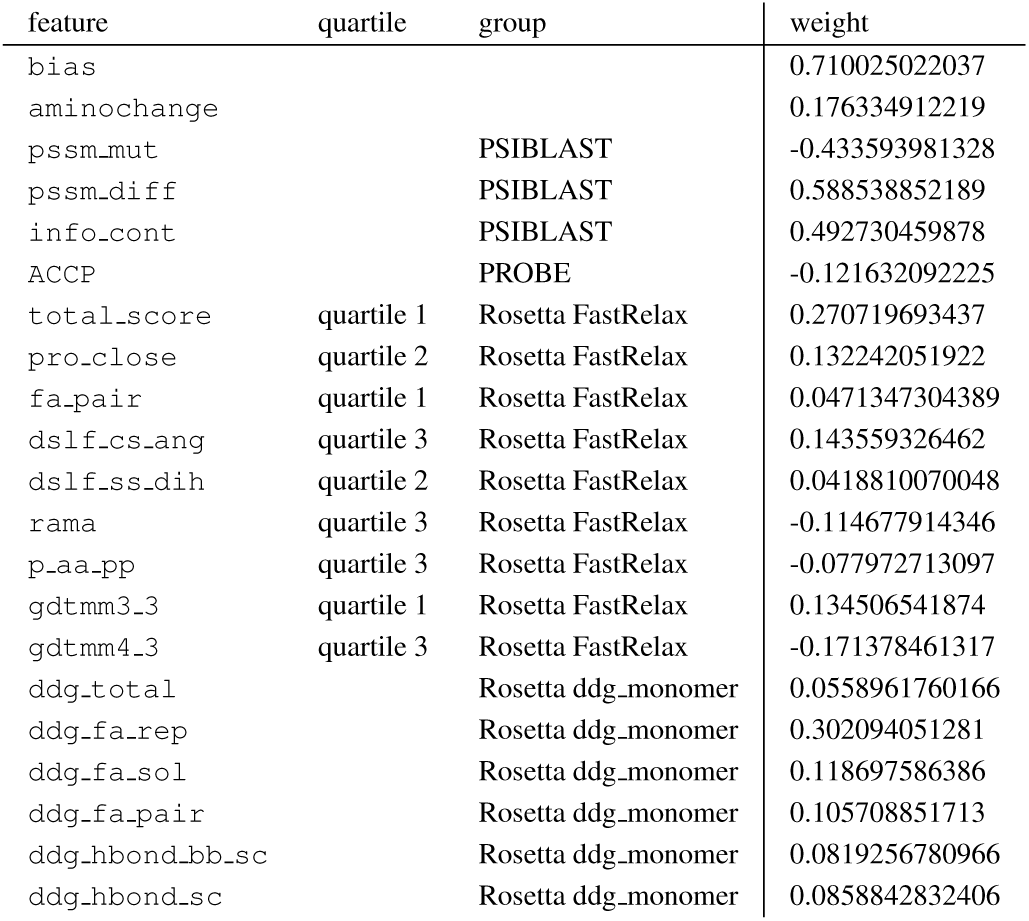
VIPUR Final Model Selected Features

*Aminochange* *Groups* We implement a crude “dissimilarity” score termed aminochange (from Poultney *et al.*(40)) by comparing the general properties of the native and variant amino acids. Each amino acid is placed into one of seven groups and a substitution is scored “1” if the native and variant amino acids belong to the same group and “2” otherwise. The amino acid groups used for aminochange are

- A, I, L, V -Alanine, Isoleucine, Leucine, Valine (small nonpolar)
- C, S, T -Cysteine, Serine, Threonine (small polar)
- D, E -Aspartic Acid, Glutamic Acid (negative charge)
- F, M, W, Y -Phenylalanine, Methionine, Tryptophan, Tyrosine (large nonpolar)
- G, P -Glycine, Proline (“bad behaved”)
- H, K, R -Histidine, Lysine, Arginine (positive charge)
- N, Q -Asparagine, Glutamine (side chain amide)

*Definitions of Surface and Buried Positions* The definition of surface and buried positions was taken from a Koga *et al.*(2012)(23), a recent publication using Rosetta for protein design. Residues were classified as “core”, “boundary”, or “surface” based on their surface area and secondary structure (reduced DSSP representation allowing helix, strand, and loop). Helix and strand residues are considered core if they have a small Solvent Accessible Surface Area (SASA≤15 Å), surface if they have a large SASA (≥60Å) and boundary if they fall between these thresholds. For loop residues, a larger SASA is tolerated for core (≤25Å) and a smaller SASA is required for surface (≥40Å). Feature distributions for the boundary residues in our training set appear very similar to the distribution for core residues, while surface residues appear distinct from both. During analysis, we considered both core and boundary classifications as “buried” or “interior” to simplify consideration of local protein environment. The VIPUR code currently assigns these positions based on a 12.5 Åcutoff (above this is considered “surface”).

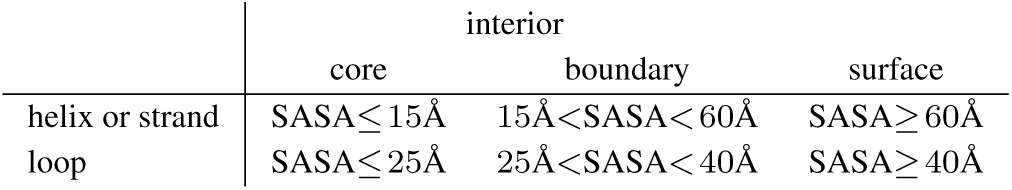

*Identification of “Essential” Positions* The structure-only classifier’s confidence metric approximates energetic destabilization, yet many variants in this dataset do not appear **deleterious** due simply to fold destabilization, likely due to the absence of interaction partners in our models. Similarly, the sequence-only classifier confidence metric serves as an appropriate approximation for amino acid conservation. These metrics correlate highly since many, but not all, destabilizing variants occur at conserved positions. Several variant positions where these scores disagree occur at interaction sites, such as W11 at the DNA binding interface of IRF1 (Supplementary Figure S10). At position 11, mutation to arginine eliminates a favorable DNA contact which “abolishes DNA binding” (UniProt annotation). Without DNA in our structural model, this mutation is not detected as destabilizing by the structure-based features even though VIPUR makes a confident **deleterious** prediction (.952), indicating conservation not caused by destabilization of the monomer structure. We observe this same behavior at other binding interfaces, such as the FOXP3 F371C dimer interface (not shown). Even though VIPUR can adequately identify disrupted interactions due to conservation of sequence or structure, incorporating these binding partners into the structural models will improve classification and enhance our automated interpretation of variant effects. Comparing the combined classifier score (destabilization vs conservation) to the structure-only classifier score (just destabilization) can identify “essential” positions that are conserved but *not* due to energetic constraints on the monomer. The precise cutoff for identifying these essential positions is unclear, though they frequently occur when the combined classifier score notably exceeds the structure-only classifier score (at interaction interfaces, score differences frequently exceed .2). The VIPUR code currently identifies potential interaction sites for **deleterious** predictions with a score difference (total-structure-only) of .2.

## SUPPLEMENTARY FIGURES

**Supplementary Figure S1.**
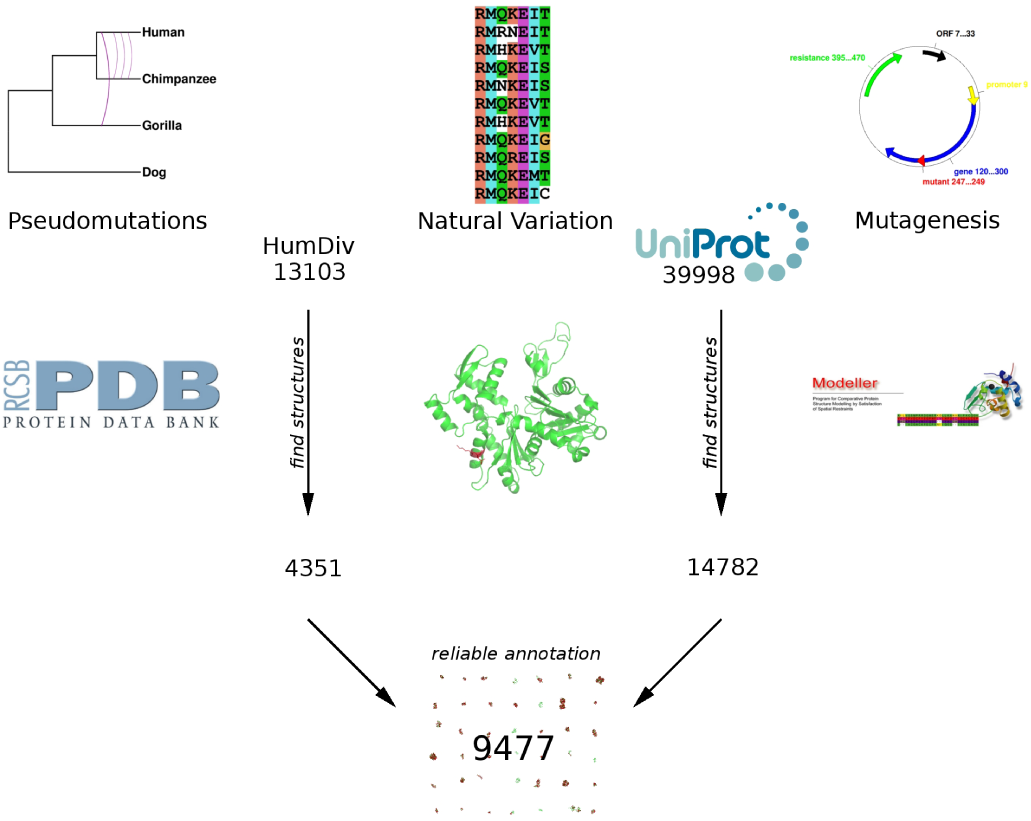
Dataset Curation: Variants with reliable annotations of deleterious or neutral effects on protein function are accumulated from UniProt and HumDiv. All variants must be mapped to structural models, limiting the number of useable annotations. Variants from all available species were extracted from UniProt reviewed entries and curated to include only reliable annotations of deleterious and neutral effects. Comparative models from ModBase and SwissModel are used when structural models are not available in the Protein Data Bank. The combined dataset reflects the diversity of variant data users are likely to use.

**Supplementary Figure S2.**
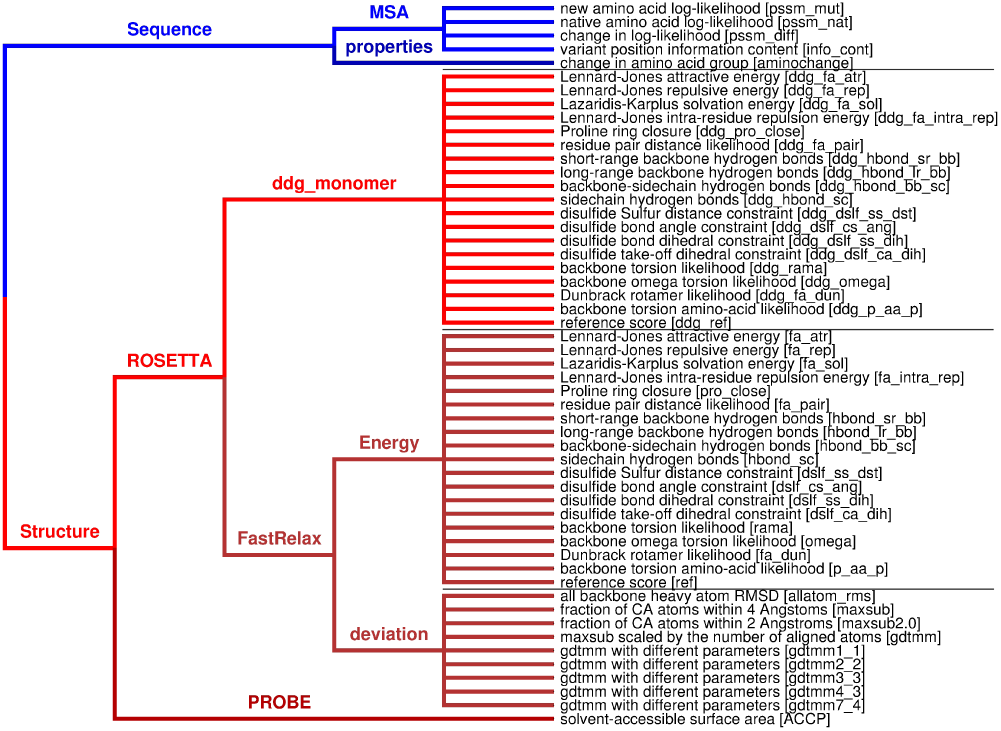
VIPUR Features. A tree represents the conceptual hierarchy of all VIPUR features. Sequence-based features (blue) are extracted directly from a PSSM output by PSIBLAST, measuring the conservation and favorability of the amino acid substitutions. Structure-based features (red) are extracted from Rosetta simulations comparing the native and variant protein structures. Variant structures are refined using the ddg_monomer protocol and the more rigorous FastRelax protocol. Every branch listed for FastRelax will become three individual features for each variant, the quartiles (Q1, Q2, Q3) of the variant score distributions evaluated on the native score distribution. Two additional features approximate the change in accessible surface area (ACCP) and changes in amino acid properties (aminochange).

**Supplementary Figure S3.**
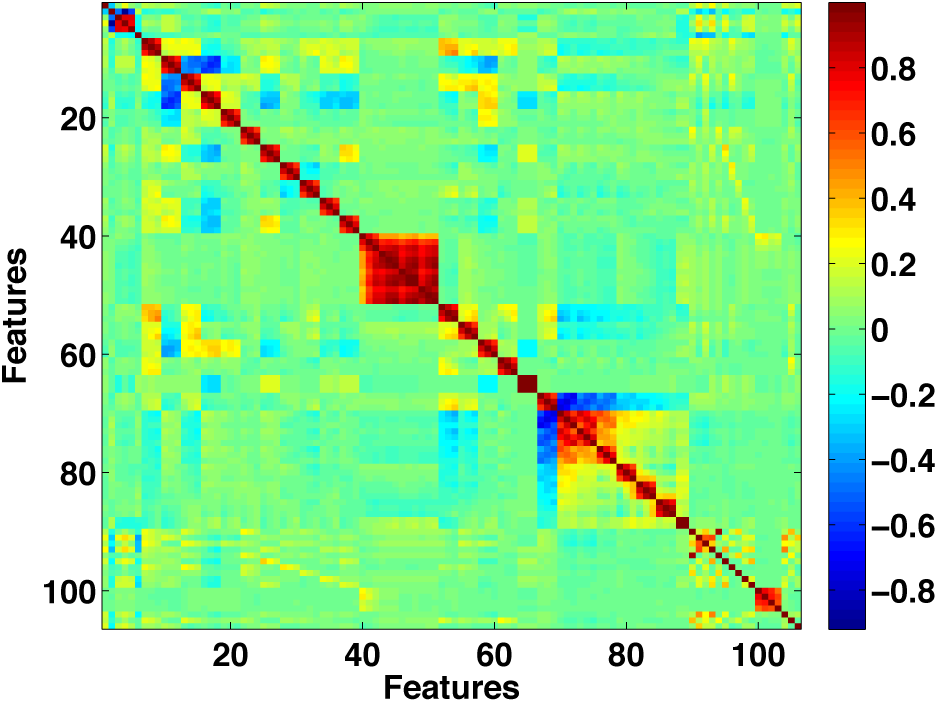
Pairwise correlation between features. Many of our features correlate highly with each other, suggesting a small subset of these features could represent the whole set with little loss. Several of these correlations are unsurprising. The three PSSM terms (features three, four, and five) contain similar information about amino acid conservation. Features derived from Rosetta FastRelax quartile analysis correlate highly with each other since three features (corresponding to the first, second, and third quartiles) are generated for each score term comparison between native and variant structures. There is also a strong correlation between the Rosetta disulfide scoring terms (blocks near the center and lower right) and gdtmm terms. Since many of these features are redundant, trained models with feature selection do not always converge on the same feature set.

**Supplementary Figure S4.**
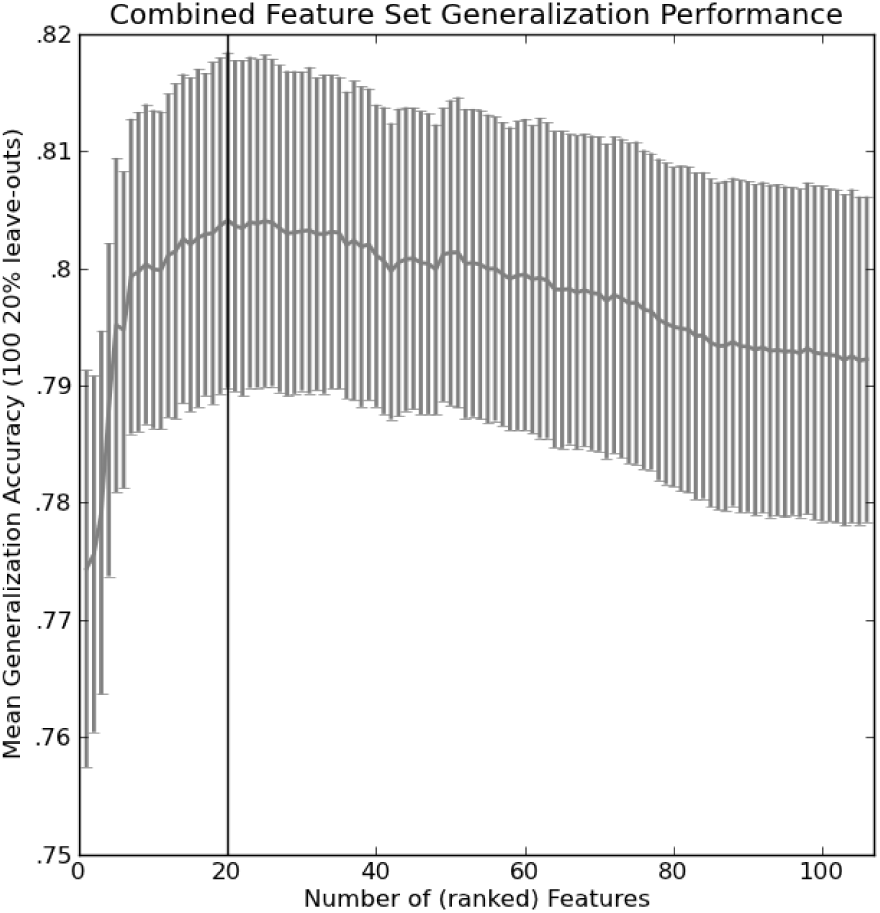
Feature Selection and Generalization. Logistic regression models were trained with the entire feature set on 100 random splits using 80% of the data (each with optimized *λ*). Features are ranked by their occurrence in these models (left, frequency of selection) and generalization performance is tested by re-training models on these ranked feature sets (100 random splits using 80% of the data for training and 20% to test generalization). Generalization performance decreases when unnecessary features are included in the model (right). The first 20 features improve performance and are used in the final model for VIPUR. Generalization decreases when additional features are added since they provide redundant information, making the model sensitive to fluctuations in the training set (overfit).

**Supplementary Table S5.** ClinVar Predictions. path-del: pathogenic variants in deleterious predictions (True Positives) at .5 cutoff, prb-neu: pathogenic variants in neutral predictions (False Negatives) at .5 cutoff, ben-del: benign variants in deleterious predictions (False Positives) at .5 cutoff, ben-neu: benign variants in neutral predictions (True Negatives) at .5 cutoff, path enrich: the ratio of path-del/ben-del, ben enrich: the ratio of path-neu/ben-neu

| method | path-del | path-neu | ben-del | ben-neu | path enrich | ben enrich |
| --- | --- | --- | --- | --- | --- | --- |
| VIPUR | 1,340 | 457 | 120 | 378 | 11.17 | 1.21 |
| PolyPhen2 | 1,590 | 207 | 177 | 321 | 8.98 | 0.64 |
| PROVEAN | 1,564 | 233 | 132 | 366 | 11.85 | 0.64 |
| SIFT | 1,509 | 288 | 148 | 350 | 10.20 | 0.82 |
| CADD | 1,587 | 210 | 126 | 372 | 12.60 | 0.56 |
| BLOSUM62 | 1,430 | 367 | 312 | 186 | 4.58 | 1.97 |

**Supplementary Figure S5.**
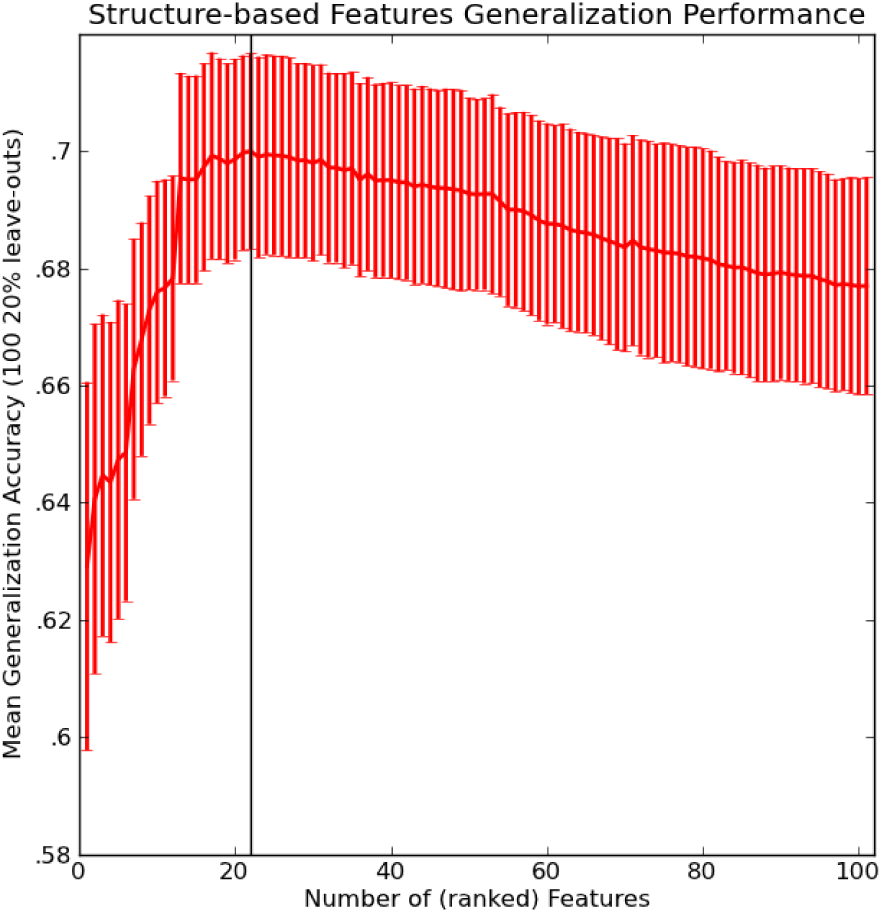
Structure-only Feature Selection and Generalization. Logistic regression models were trained with just the structure-based features on 100 random splits using 80% of the data (each with optimized *λ*). Structure-based features are ranked by their occurrence in these models (left, frequency of selection) and generalization performance is tested by re-training models on these ranked feature sets (100 random splits using 80% of the data for training and 20% to test generalization). Generalization performance decreases when unnecessary features are included in the model (right). For the structure-based features, the first 22 features improve performance. Generalization decreases when additional features are added since they provide redundant information, making the model sensitive to fluctuations in the training set (overfit).

**Supplementary Figure S6.**
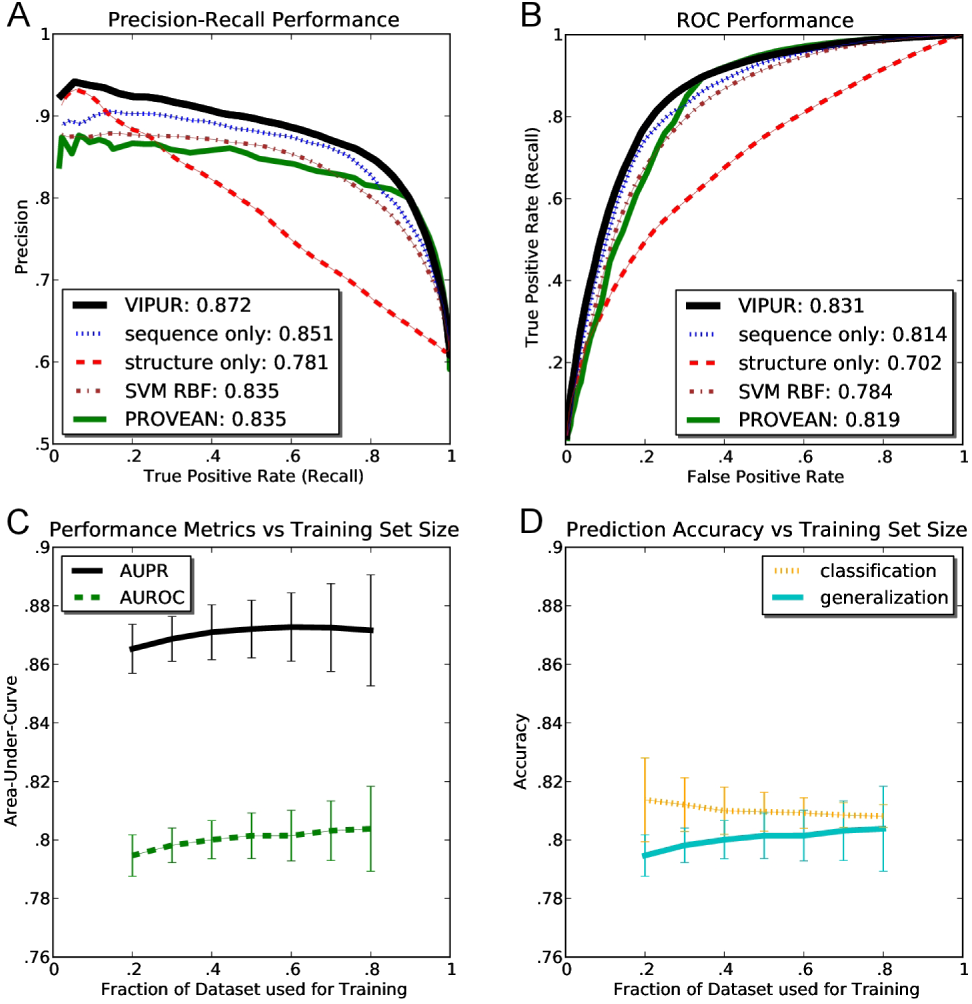
Precision-Recall and ROC curves. Averaged from models trained on 100 random dataset splits (80% training, 20% testing). Performance of PROVEAN (green) and a Radial Basis Function Support Vector Machine (brown) classifier are shown for comparison. A) VIPUR achieves a higher AUPR than other methods due to the enhanced precision of structure-based features. Sequence based methods (PROVEAN and our sequence-only classifier) generalize well, but cannot indicate confident predictions. VIPUR’s confidence score scales with precision, clearly indicating which predictions are more likely to be correct. B) ROC performance similarly shows that VIPUR is more specific than other methods at high confidence scores. PROVEAN and our sequence-only classifier, clearly indicate this sensitivity-specificity tradeoff. C) These model properties are robustly produced during training and do not change much when trained on more than 50% of our training set. D) Classification and generalization (leave-out) performance converge for our feature set, indicating our model is not overfit and performance estimates are reliable.

**Supplementary Figure S7.**
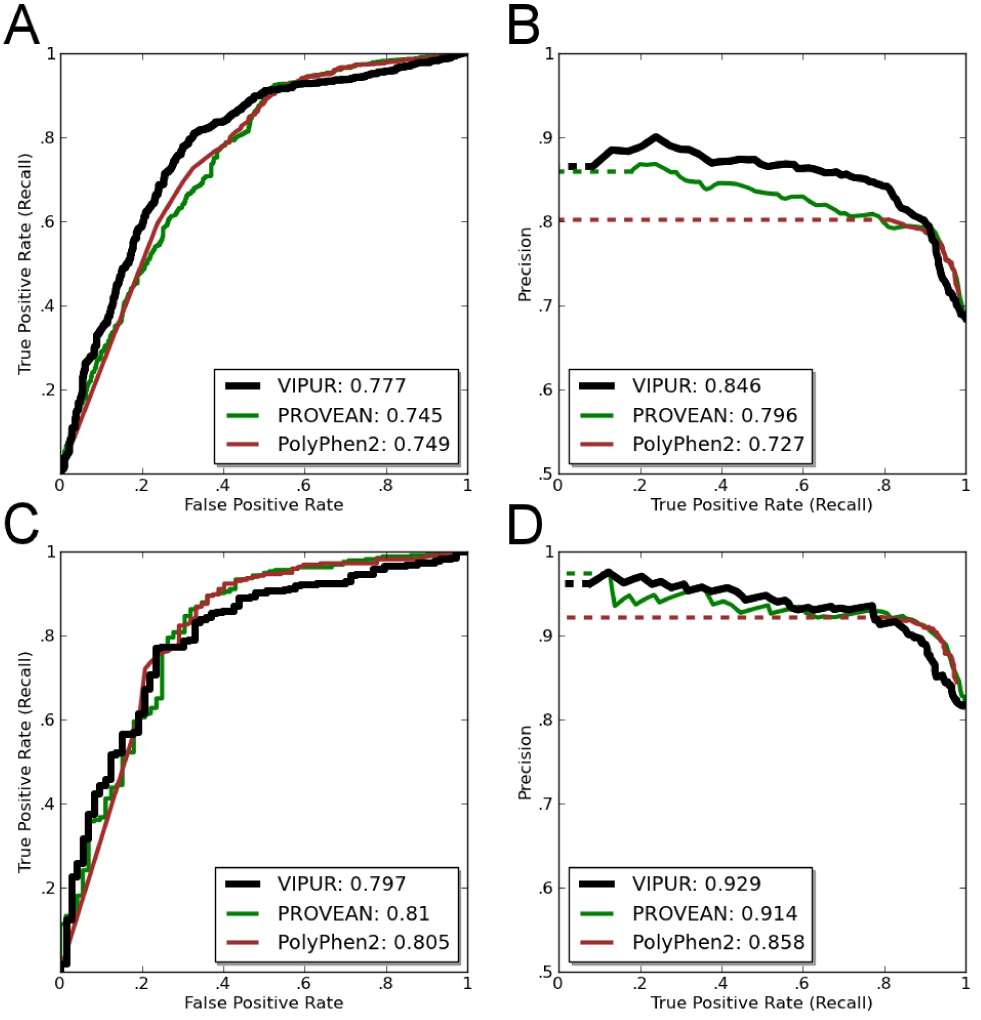
Precision-Recall and ROC curves on a subset of human variants. AUROC and AUPR curves are calculated for a set of human variants in VTS. These variants are not contained within HumDiv and A VIPUR classifier is retrained on our training set excluding these variants, allowing the VIPUR feature space to be compared with PolyPhen2. A) VIPUR has a slightly increased AUROC compared to PROVEAN and PolyPhen2, but notably higher AUPR (B, C), indicating VIPUR’s top predictions are more enriched for true deleterious variants compared to other methods. D, E, F) PROVEAN and PolyPhen2 have notably improved performance on common human variants, suggesting these methods may be overfit to this type of variation.

**Supplementary Figure S8.**
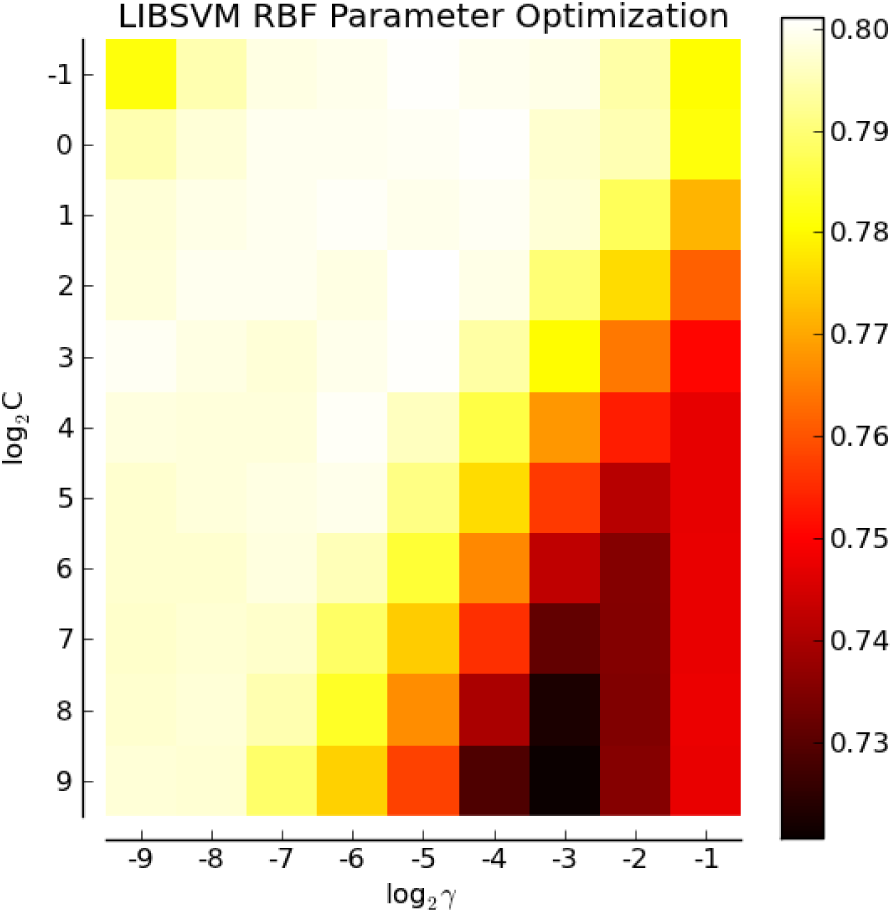
Parameter optimization for a Radial-Basis Function Support Vector Machine using LibSVM. Optimum values for the meta parameters C(SVM cost) and γ(RBF variance) were chosen by training over *C*∈{2^−1^,2^0^,2^1^,&,2^9^}and *γ*∈{2^−8^,2^−7^,&,2^−1^}. The values (C=2^2^, *γ*=2^−4^) maximize accuracy using 5-fold cross-validation (highest average accuracy on the leave-out sets).

**Supplementary Figure S9.**
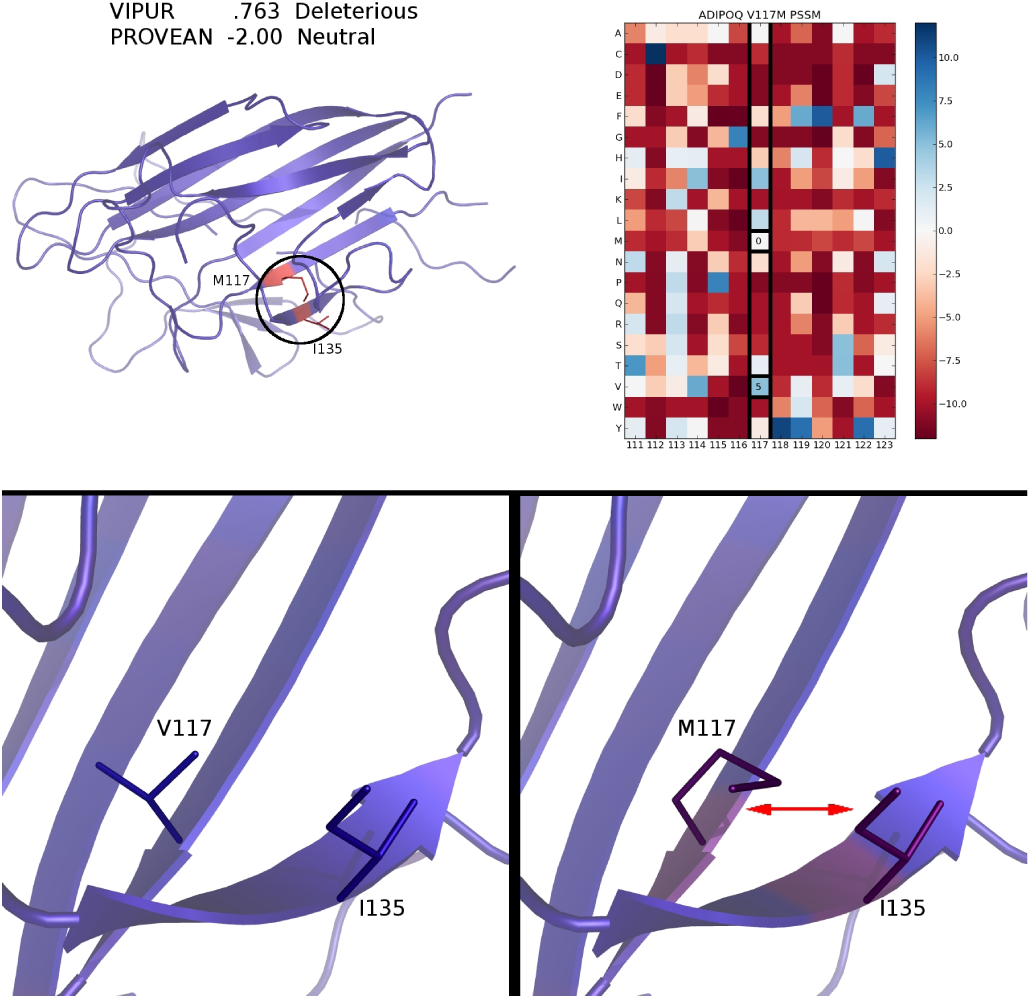
V117M disrupts strand pairing for ADIPOQ. V117M is predicted deleterious (.763) due to statistically unfavorable backbone conformation and predicted neutral (-2.00) by PROVEAN. Every residue in ADIPOQ is colored by the difference in Rosetta energy between the native and variant protein structures, highlighting the destabilization introduced by V117M (top left). The PSSM generated by PSIBLAST does not indicate strong conservation at position 117 (top right, PSSM columns shown for surrounding residues). The native V117 structure forms stable hydrophobic contacts and strand pairing (bottom left, residues colored by Rosetta energy of a representative model). The M117 variant model cannot accommodate the larger amino acid without disrupting strand pairing.

**Supplementary Figure S10.**
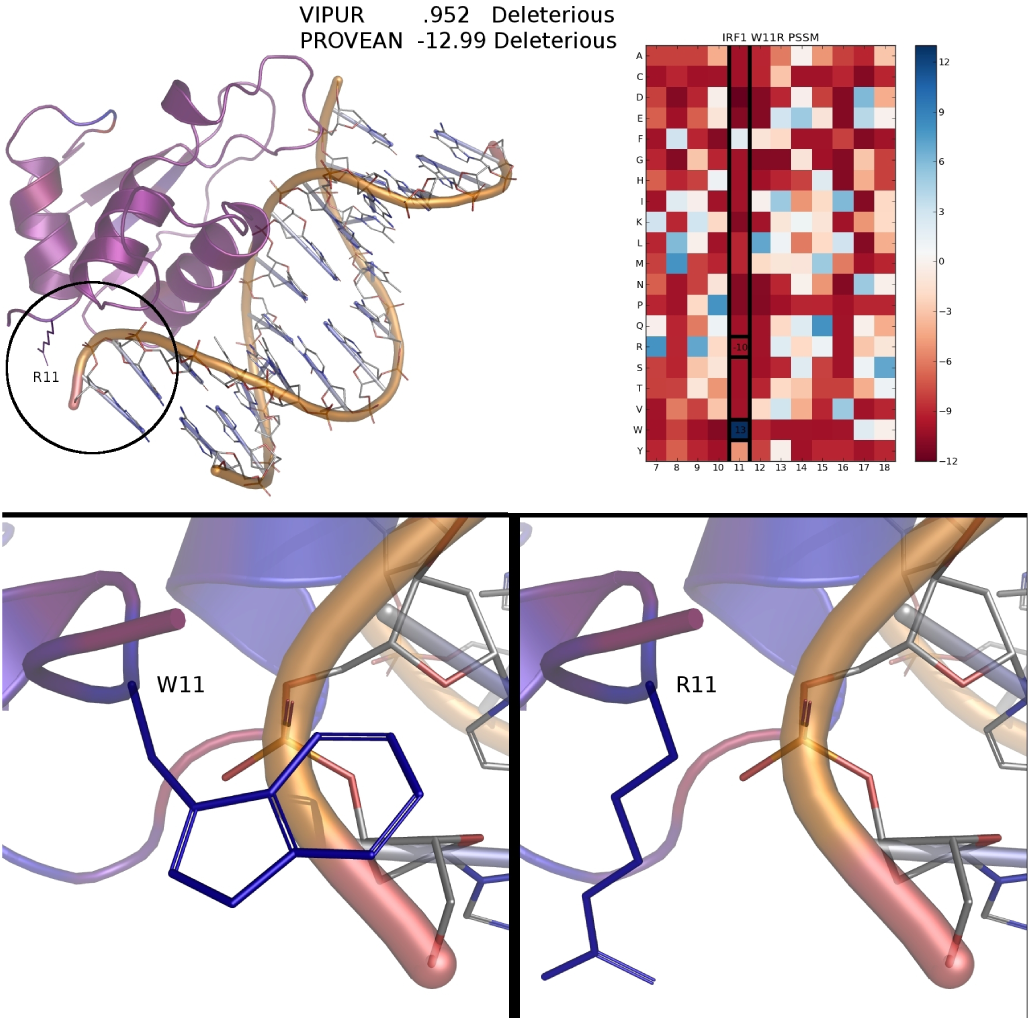
W11R eliminates a DNA contact. VIPUR predicts W11R is deleterious (.952), matching the UniProt annotation that this variant “abolishes DNA binding”. Our structural model lacks DNA and the correct prediction is due to the sequence-based features. Every residue in IRF1 is colored by the difference in Rosetta energy between the native and variant protein structures, demonstrating the similarity of energies between native and variant structures (top left). The PSSM generated by PSIBLAST indicates that W11 is highly conserved (top right, PSSM columns shown for surrounding residues). W11 packs against the DNA backbone in the template PDB 1IF1 (bottom left, residues colored by Rosetta energy of a representative model) and this contact is lost in R11 (bottom right). Without DNA in the structural model, W11R is not detected as energetically destabilizing, however, the sequence-based features accurately inform a deleterious prediction.

**Supplementary Figure S11.**
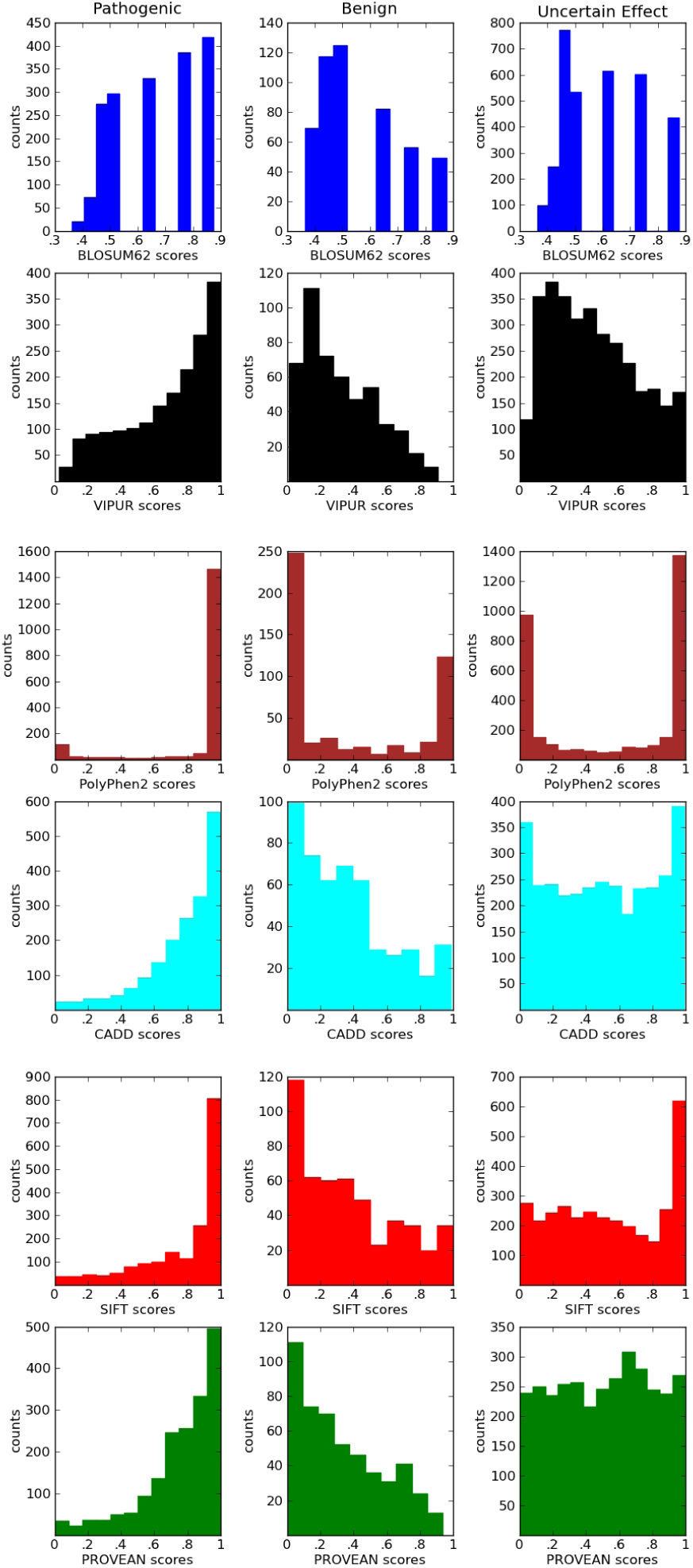
ClinVar Predictions. Score histograms for several methods on ClinVar pathogenic, benign, and uncertain variants. All methods match expectations for pathogenic and benign variants, with pathogenic variants having a skewed distribution of deleterious scores and benign variants having a broad distribution of neutral scores. PolyPhen2 predictions are notably pushed to high and low values. Predictions on variants with uncertain labels are very diverse and suggest very different error rates between available methods.

**Supplementary Figure S12.**
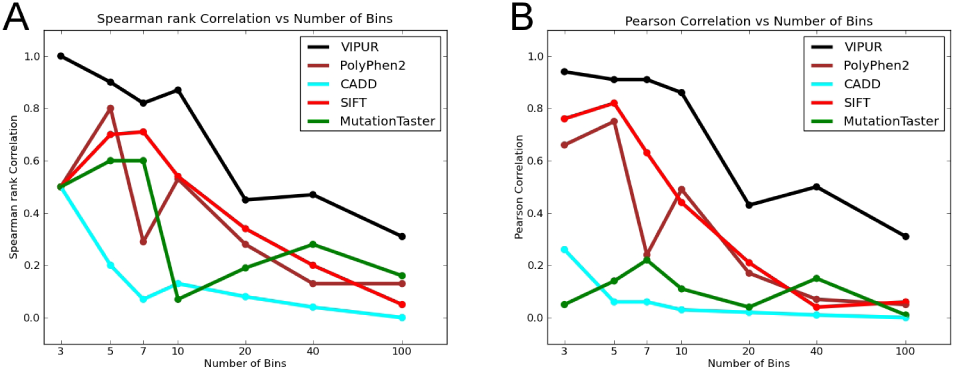
Simons Simplex Collection Proband Enrichment is Robust to the Bin Size. Altering the number of bins used for the correlation calculation between deleterious scores and proband enrichment changes the correlation value obtained but not the trend. These methods predict categorical labels from a continuous deleterious score where low values indicate neutral mutations and high values indicate highly disruptive mutations. VIPUR predictions produce higher correlation values than other pathogenicity prediction methods independent of the number of bins used for the calculation.

**Supplementary Figure S13.**
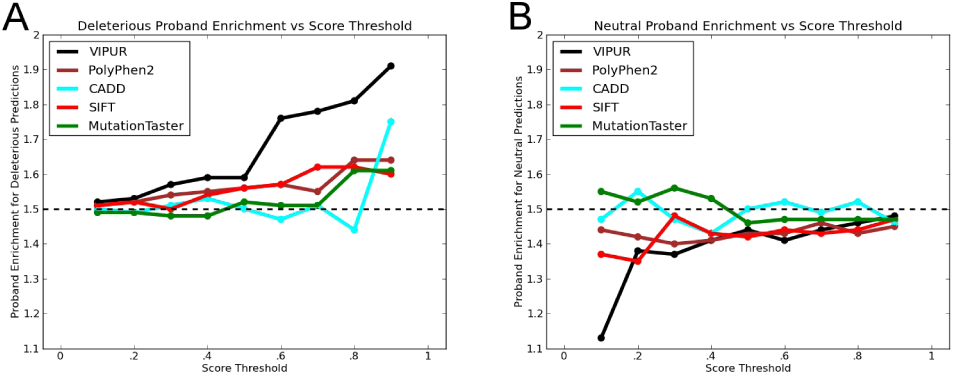
Simons Simplex Collection Analysis Proband Enrichment is Robust to the Score Threshold. Altering the threshold used to determine deleterious vs neutral predictions changes the enrichment ratio obtained but does not notably alter the trend. This evaluation is the accumulated version (accumulated to 1 for deleterious predictions and to 0 for neutral predictions) of the bin correlation (Figure 6). VIPUR predictions are consistently above the background ratio for deleterious predictions and below the background ratio for neutral predictions, with sharp increases for high confidence predictions. PolyPhen2 and SIFT both maintain the appropriate trends but are have consistently worse enrichment ratios compared to VIPUR.

